# Single-cell RNA Sequencing Reveals Methamphetamine Inhibits the Liver Immune Response with Involvement of the Dopamine D1 Receptor

**DOI:** 10.1101/2024.06.26.600774

**Authors:** Jin-Ting Zhou, Yungang Xu, Xiao-Huan Liu, Cheng Cheng, Jing-Na Fan, Xiaoming Li, Jun Yu, Shengbin Li

## Abstract

Methamphetamine (METH) is a highly addictive psychostimulant that causes physical and psychological damage and immune system disorder, especially in the liver, which contains a significant number of immune cells. Dopamine, which is a key neurotransmitter in METH addiction and immune regulation, plays a crucial role in this process. In this study, we developed a chronic METH administration model and conducted single-cell RNA sequencing to investigate the effect of METH on liver immune cells and the involvement of the dopamine receptor D1 (DRD1) in this process. Our findings revealed that chronic exposure to METH induced an immune cell shift from Ifitm3+Mac and Ccl5^+^Mac to Cd14^+^Mac, and from Fyn^+^CD4^+^Teff, CD8^+^T, and NKT to Fos^+^CD4^+^T and Rora^+^ILC2, along with suppression of multiple immune functional pathways. DRD1 was implicated in the regulation of some of these pathways and the shifts of hepatic immune cells. This research provides valuable insights into the development of therapies aimed at mitigating METH-induced immune impairment.

## Introduction

Methamphetamine (METH or MA, also known as ice) is a highly addictive and neurotoxic drug, affecting over 36 million people globally in 2021 [1, 2]. METH causes severe societal problems and harm to health, damaging the central nervous system and immune system, leading to susceptibility to infectious diseases like viral hepatitis and HIV [3–6]. Despite increasing studies, the exact mechanisms underlying the effect of METH on the immune system remain unclear.

Chronic METH use disrupts immune cell balance [4, 7, 8], affecting signaling pathways and causing damage to cells like splenic dendritic cells (DCs) and T cells [4]. Chronic METH use also influences various pathways such as chemokine receptors and intracellular calcium levels, all of which affect immune responses [4]. The liver, which is a frontline immune organ [6, 9], experiences METH-induced toxicity [10], leading to hepatotoxicity, cell cycle arrest [11, 12], and reduced clearance [13]. However, there are limited studies exploring changes in liver immune cells due to chronic METH use.

The dopamine (DA) system is crucial in METH-related disorders. DA facilitates neurotransmission through the activation of five DA receptors: D1-like DA receptors (DA receptor D1, DRD1; DA receptor D5, DRD5), D2-like DA receptors (DA receptor D2, DRD2; DA receptor D3, DRD3), and DA receptor D4 (DRD4) [14]. METH administration markedly increases DA levels in the nucleus accumbens [15]. Extracellular DA is crucial for activating DA receptors and interacts with DRD1, DRD2, and DRD3 in the nucleus accumbens [14]. The rewarding properties of METH rely on the presence of DRD1 and DRD2 [16, 17], and METH as a major addictive psychostimulant through reward circuits, led to physical and psychological alterations [15, 18]. METH causes dopamine (DA) to be released extracellularly and reduces dopamine reuptake through various mechanisms, such as blocking the role of dopamine transporter (DAT) [19] and increasing the expression of DRD1 [17]. A DRD1 antagonist has been reported to extinguish METH-induced conditioned place preference [20, 21]. In addition to being a neurotransmitter, DA is an important regulator of immune function [22]. Immune cells produce DA, which can be used as an autocrine/paracrine mediator of the immune cells themselves and adjacent cells [23]. DA receptors and other DA-related proteins were detected in many immune cells, suggesting that DA is a crucial component in regulating immune function [24, 25]. However, whether METH regulates liver immunity, and the underlying mechanisms of METH in regulating immune function remain elusive. Therefore, investigating the regulatory mechanisms of METH through DA is a key step to understanding the impact of METH on liver immunity.

Single-cell RNA sequencing (scRNA-Seq) offers more powerful analyses with higher resolution compared with conventional bulk RNA sequencing (RNA-Seq). Using scRNA-Seq, cell subsets, subset functions, and cell–cell interactions in complex tissues can be identified at the single-cell transcriptomic level. METH has been reported to induce hepatotoxicity and cause deleterious inflammatory responses [26, 27]. However, using scRNA-Seq analysis to investigate the effects of METH on liver immune cells has not been reported. Therefore, in this study we used scRNA-Seq to explore METH-induced immunosuppression in the liver through DRD1. Examination of liver tissues from DRD1 knockout and wild-type METH-chronic treatment mice revealed changes in immune cell numbers and functions. This research might offer insights into treating METH addiction by understanding the impact of this drug on liver immunity.

## Results

### METH exposure and DRD1 deletion caused changes of immune cells in mouse liver revealed by RNA-Seq

To explore the effects of METH on hepatic immune cells and whether DRD1 was involved, bulk RNA-Seq was performed with liver tissues from the WS, DS, WM, and DM mice. Enrichment analysis of the DEGs among the showed that many immune-related pathways were affected by METH and DRD1 knockout, including response to interferon, defense response to virus, and regulation of innate immune response (Figure S1A; Tables S1, 2). In addition, immune infiltration analysis showed changes in the proportion of some liver immune cells caused by METH exposure as well as DRD1 deletion. Flow cytometry analysis further supported that METH exposure and DRD1 deletion altered the proportion of certain cells; for example, the WM group had increased CD4T cells and decreased CD8T cells, whereas the DM group had a high proportion of CD8T cells and a trend of decreased macrophages (Figure S1B, C). The results suggested that METH exposure affects the number and function of liver immune cells, with DRD1 involved in the process. Consequently, we further investigated the target subclusters of liver immune cells to explore the mechanisms of METH and possible involvement of DRD1.

### ScRNA-Seq maps distinct immune cell populations

After quality control and filtering, we obtained 21743 transcriptomes of single cells from hepatic NPCs isolated from the three groups of mice (*n* = 3 from the WS group, *n* = 4 from the WM and DM groups) **(Figure 1A)**. Based on the expression of canonical gene markers, 13 distinct cell clusters were identified **(Figure 1B, C)**. In total, we identified nine major immune cell clusters with their unique markers (**Figure 1D**; Table S3), including macrophages (Mac, *Lyz2*^+^), plasmacytoid dendritic cells (pDC, *Siglech*^+^), T cells (T, *Cd3d*^+^), natural killer cells (NK, *Nkg7*^+^), granulocytes (Gran, *Csf3r*^+^), dendritic cells (DC, *Bst2*^+^), plasma B cells (Plasma-B, *Jchain*^+^), neutrophils (Neut, *Camp*^+^), and B cells (B, *Cd79a*^+^) **(Figure 1E)** [28, 29]. A cluster identified with highly expressed Mki67 (markers for cell proliferation) [28, 30] was named dividing cells (4_Dividing). This cluster contains NK and T cells, myeloid cells, and hepatocytes (**Figure 1D**, Figure S2). These results indicated that the 4_Dividing cluster may be the progenitors of the aforementioned cells.

**Figure 1.**
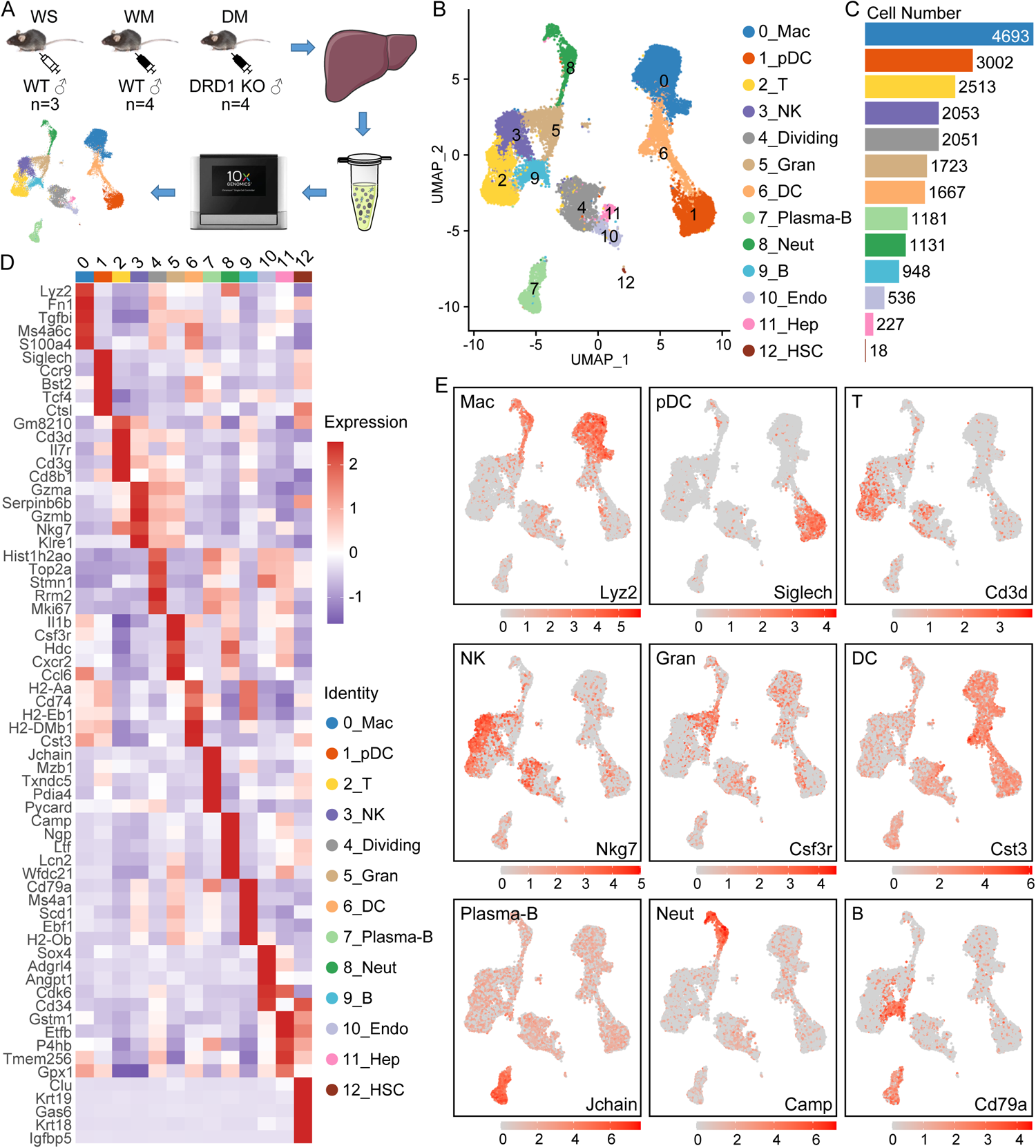
Single-Cell RNA-Seq reveals immune cell populations in mice liver. **A.** Schematic diagram for single-cell transcriptomic profiling of mice NPCs. WS: WT + Saline; WM: WT + METH; DM: DRD1 knockout + METH. **B. & C.** The 13 types of hepatic nonparenchymal cells and their counts. **D.** The signature genes for each cell type. **E.** The expression profiles of representative marker genes of main immune cell types.

### METH induces immunosuppressive hepatic microenvironments partially through DRD1

To investigate the distinctive immune profiles of different groups, we analyzed the composition of immune cells in each group (**Figure 2A, B**; Table S4). We found that macrophages were enriched in the WS and WM groups, and neutrophils in the WM group. In the DM group, more than 50% of T cells were preferentially enriched, but macrophages and myeloid cells decreased significantly, indicating that DRD1 was required for homeostasis of hepatic myeloid cells, and the opposite effect of DRD1 between T cells and the myeloid cells.

**Figure 2.**
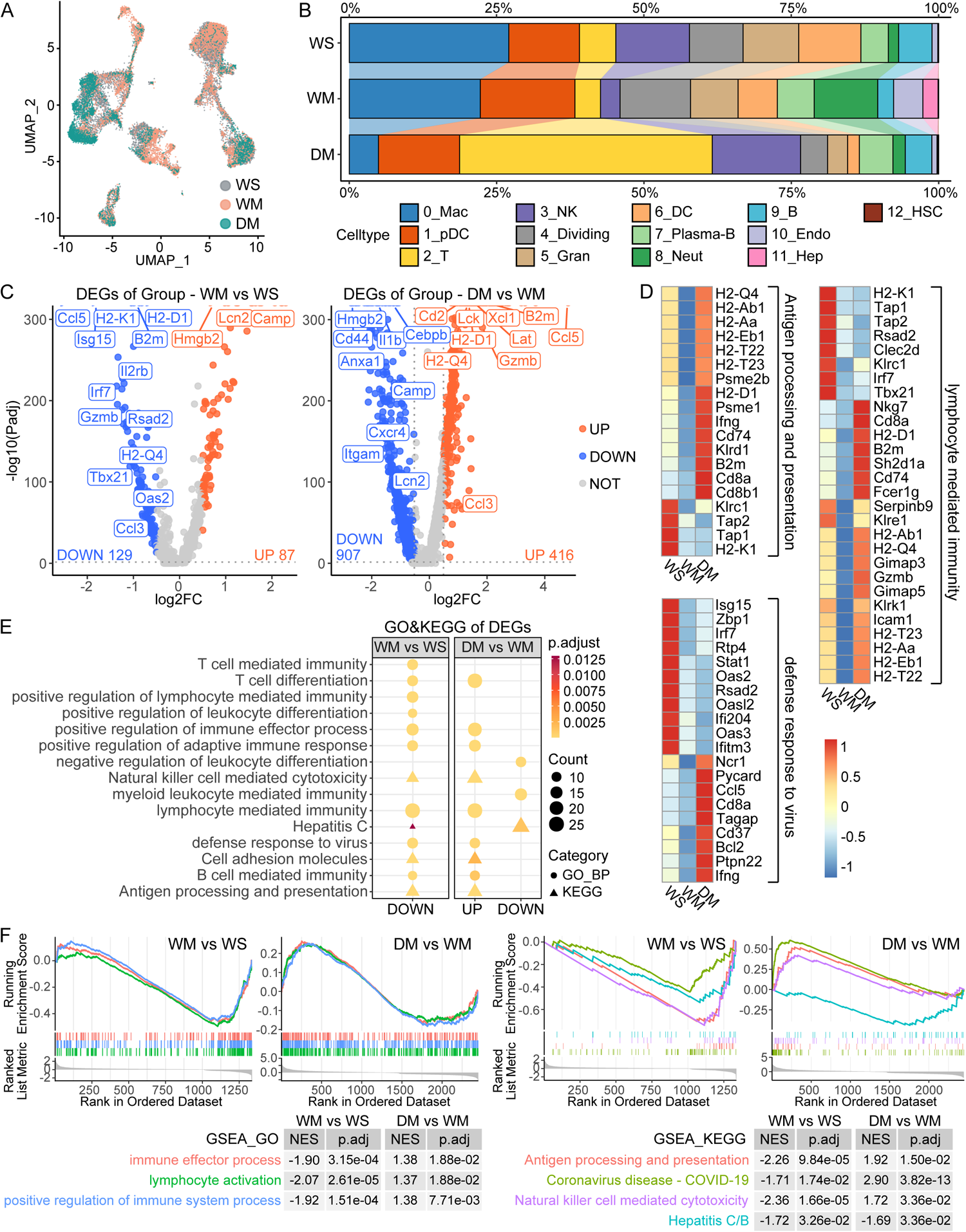
Comparative transcriptomic analyses reveal the METH-induced immunosuppressive hepatic microenvironments. **A.** The cell populations from 3 groups of mice. **B.** The cell proportion of each cluster in 3 groups. **C.** Volcano plots showing fold change of gene expression (log2 scale) for down-regulated and up-regulated genes of WM vs. WS and DM vs. WM. Up-regulated genes (P.adj < 0.05 and log2FC > 0.5) are shown with red dots, down-regulated genes (P.adj < 0.05 and log2FC < −0.5) are shown with blue dots, and insignificant genes (P.adj > 0.05 or log2FC < 0.5) are shown with gray dots. **D.** The relative gene expressions of the representative immune pathways. **E.** GO and KEGG enrichment for the up-regulated and down-regulated genes in (C). **F.** GSEA enrichment of all DEGs (P.adj < 0.05).

Differential gene expression analysis revealed 87 up-regulated and 129 down-regulated genes between the WM and WS groups, and 416 up-regulated and 907 down-regulated genes (│log2FC│> 0.5) between the DM and WM groups (**Figure 2C**; Table S5). The enrichment analysis of the DEGs between the WM and WS groups showed the down-regulation of abundant immune-associated functional pathways, including antigen processing and presentation, lymphocyte-mediated immunity, positive regulation of immune effector process, NK cell-mediated immunity, defense response to virus, B cell-mediated immunity, and cell adhesion molecules. However, when comparing the DM and WM groups, these pathways were up-regulated (**Figure 2D–F**; Tables S6, 7). These findings indicate that chronic exposure to METH induces immunosuppressive hepatic microenvironments and that DRD1 knockout could partially reverse the effect. In the gene lists of the above pathways, a few DEGs were down-regulated in the WM group but up-regulated in the DM group, including H2 (histocompatibility-2) genes (encoding subunits of MHC I and MHC II), *Cd2*, *Icam1*, *Zap70*, *Il2rg*, *Gimap3/5* (GTPase IMAP family member), *Serpinb9*, *Irf7*, *Gzmb*, *Ifng*, and *Cd74* (**Figure 2D**, Figure S3), suggesting these genes might be suppressed by METH through DRD1.

### METH exposure increases immunosuppressive macrophages but DRD1 deletion causes macrophage loss

As the biggest immune cell population, 4693 macrophages were identified **(Figure 3A)**. Six subclusters were further annotated, including c0_Mac-Ifitm3, c1_Mac-Cd14, c3_Kupffer-Lgmn, c2_Mac-Ccl5, c4_Mac-Adgre4, and c5_Mac-Il1b (**Figure 3A, B**; Table S8). We observed significant differences in the compositions of these six subclusters between the three groups of mice. The proportions of c0_Mac-Ifitm3 and c2_Mac-Ccl5 in the WS group were significantly higher than those in the WM and DM groups, c1_Mac-Cd14 and c4_Mac-Adgre4 were higher in the WM group, and the proportion of Kupffer cells was higher in the DM group (**Figure 3C, D**).

**Figure 3.**
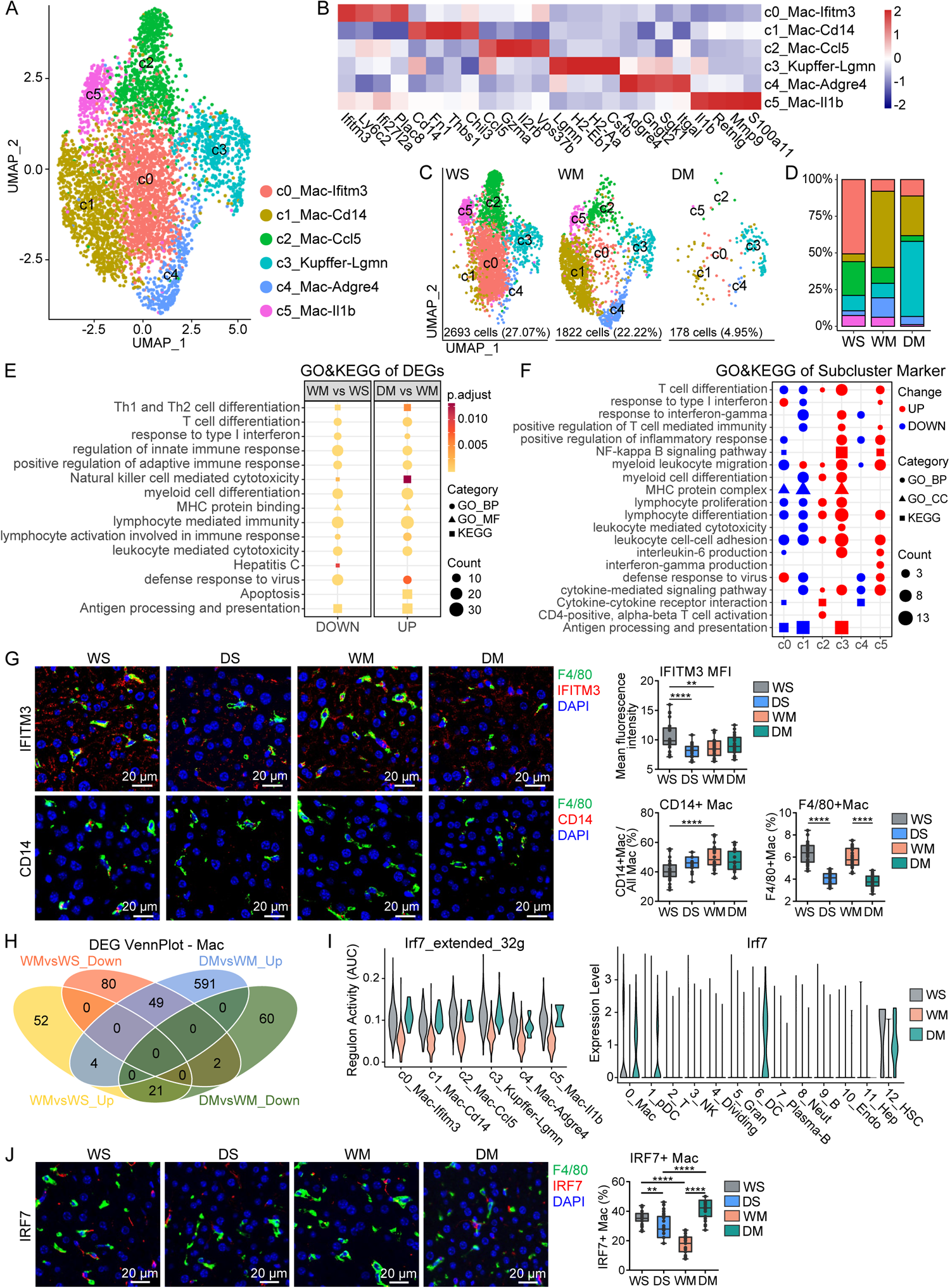
METH increase immunosuppressive macrophages. **A.** Six subclusters were annotated from the macrophages. **B.** The expression of marker genes for each macrophage subcluster. **C. & D.** Group-wise cell populations and proportions of macrophage. The legend is shared with (A). **E.** GO and KEGG enrichment of the up-regulated and down-regulated genes. **F.** GO and KEGG enrichment of the marker genes from the subclusters of macrophages. **G.** Representative double immunofluorescent of mice hepatic tissue (n=3) with DAPI nuclear counterstain (×40, bar=20 μm) (left). Quantification of mean fluorescence intensity of IFITM3, CD14+Macrophages and Macrophages (right), **: P < 0.01, ****: P < 0.0001. **H.** The intersection of up-regulated and down-regulated genes of Macrophages. **I.** Regulon activity (AUC) of Irf7 and its targets (left) and expression level of Irf7 in all cell types (right). **J.** Representative double immunofluorescent of mice hepatic tissue (n=3) with DAPI nuclear counterstain (×40, bar = 20 μm) (left). Quantification of IRF7+Macrophages (right), **: P < 0.01, ****: P < 0.0001.

Compared with the WS group, the proportion of macrophages in the WM group decreased slightly. Unexpectedly, compared with the WM group, the proportion of macrophages in the DM group decreased markedly and almost disappeared (**Figure 3C**). Furthermore, cell cycle scores showed a G_2_M cell cycle arrest in the WM group, and a G_1_ cell cycle arrest in the DM group, which may suggest a suppression of proliferation in these two groups (Figure S4A). The DEGs analysis showed that there were more down-regulated genes in the WM group compared with the WS group, and these genes were enriched in many immune function pathways, for instance, antigen processing and presentation, defense response to virus, and T cell differentiation. Most of these pathways were reversed in the DM group, indicating that the impaired immunocompetence caused by METH exposure may occur via a mechanism involving DRD1 (**Figure 3E**; Figure S4B; Tables S9, 10).

To identify functional changes of the shifts from c0_Mac-Ifitm3 and c2_Mac-Ccl5 in the WS group to c1_Mac-Cd14 caused by METH exposure in the WM group, GO and KEGG enrichment analyses of the marker genes were performed (**Figure 3F**, Table S11). Compared with c0_Mac-Ifitm3, which exhibited up-regulation of the defense response to virus pathway, c1_Mac-Cd14 had down-regulated myeloid leukocyte differentiation, leukocyte-mediated cytotoxicity, and defense response to virus.

Subcluster c2_Mac-Ccl5 uniquely showed up-regulated lymphocyte proliferation and CD4^+^αδ T cell activation pathways. Subcluster c3_Kupffer-Lgmn showed enhanced immune-related functions and may be functionally active. Subcluster c4_Mac-Adgre4 was only enriched with a relatively small number of down-regulated pathways, but contained highly expressed genes such as *Adgre4*, *Gngt2*, and *Sgk1*, which may be related to intercellular information interactions. Subcluster c5_Mac-Il1b uniquely up-regulated the Ifng production pathway, indicating its immune functional activation. The percentages of c0_Mac-Ifitm3 and c2_Mac-Ccl5 were both decreased and shifted into c1_Mac-Cd14 in the WM group, again demonstrating that METH exposure leads to suppression of immune functions **(Figure 3D, G)**. In the DM group, c3_Kupffer-Lgmn was significantly increased, which may not fully compensate for the suppressed function of macrophages because their number was markedly reduced. The change in the proportion of Kupffer cells between the WM and WS groups is similar to macrophages, but different in the DM group, in which the proportion was increased **(Figure 3D)**. The DEGs and the enrichment findings of Kupffer cells were slightly different from those of macrophages (Figure S4C–E; Tables S12, 13).

Pathways such as antigen processing and presentation, T cell-mediated immunity, response to type I interferon, and defense response to virus were down-regulated in the WM group and up-regulated in the DM group, indicating that METH exposure suppresses Kupffer immune function but DRD1 deletion partially reverses this suppression. However, some pathways, such as myeloid leukocyte migration and leukocyte chemotaxis were up-regulated in both comparison groups.

Through Venn diagram analysis (VDA), we identified some genes regulated by METH that were regulated reversely by DRD1 deletion, including membrane protein *Ifitm3*, MHC, and transcription factor *Irf7* (**Figure 3H**; **Table 1**; Table S14). An interaction network of these genes based on the STRING Database was generated (Figure S4F). We further performed transcription factor analysis by SCENIC on 0_Mac and found that the expression of *Irf7* and its regulatory network (regulon activity) were down-regulated in immune-suppressed subclusters c1_Mac-Cd14 and c4_Mac-Il1b, which consisted of most macrophages in the WM group (**Figure 3I**; Table S15). Decreasing *Irf7*, which is a key transcriptional regulator of type I interferon (IFN)-dependent immune responses, demonstrates the inhibitory effect of METH on macrophage immune responses [31]. The expression changes of *Irf7* in macrophages of the liver were verified by tissue immunofluorescence **(Figure 3J)**.

**Table 1.**
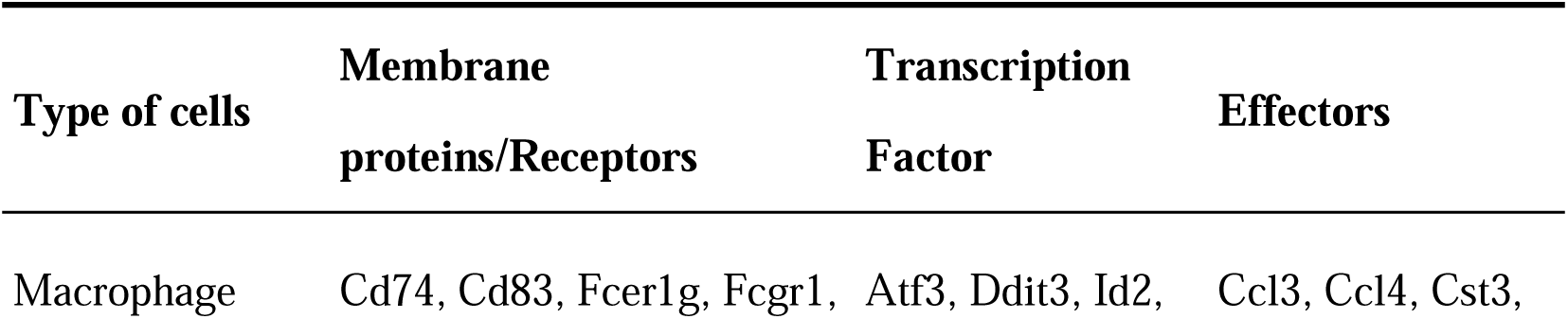
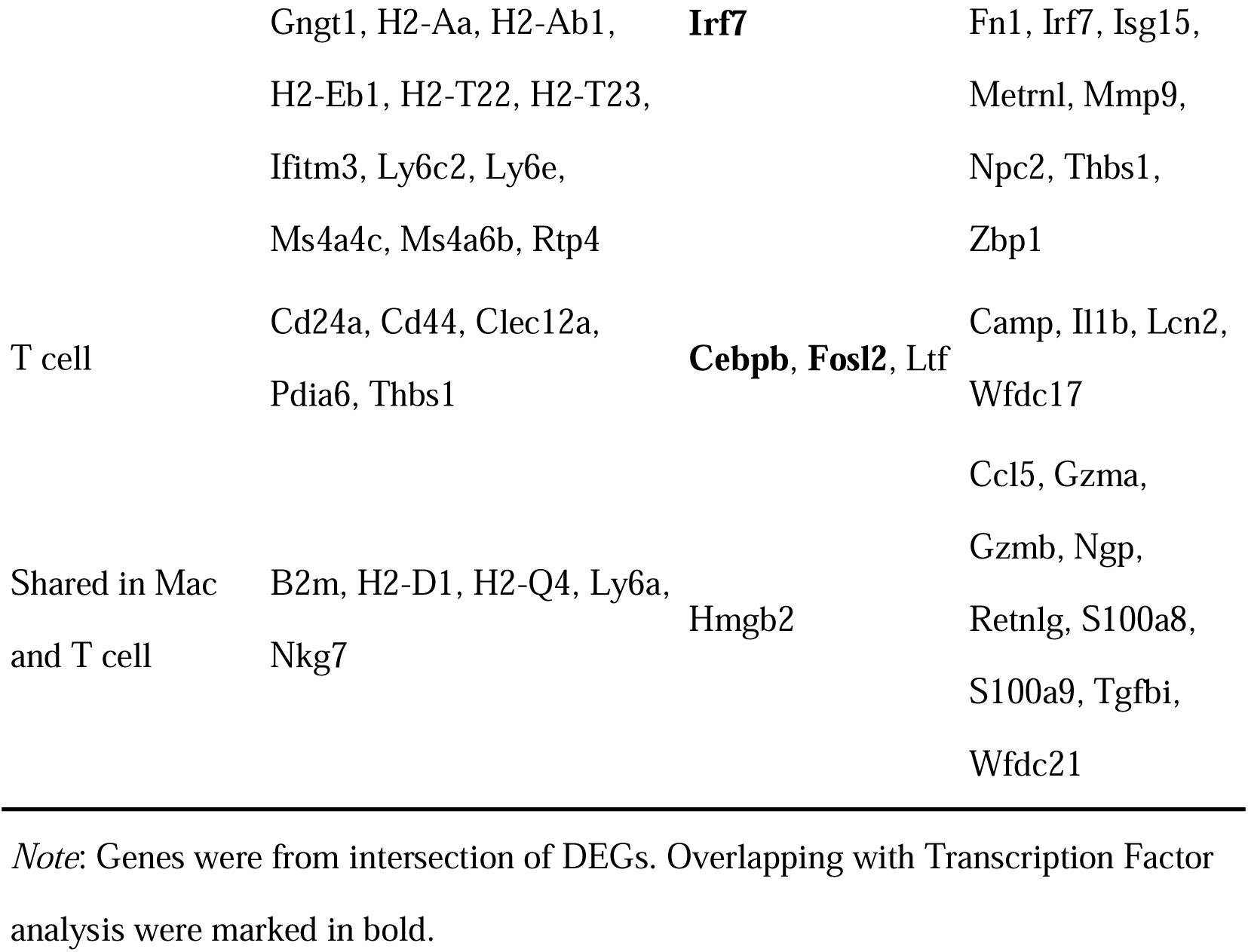
Genes regulated by METH through DRD1.

### METH exposure induces T cell immunodeficiency but DRD1 knockout partially prevents this activity

A total of 2513 T cells were identified according to traditional immunological surface markers (Figure 4A). The T cells were divided into 10 subclusters, including four clusters for CD8^+^T cells: c2_Cd8-Klk8-Tcm (*Cd8^+^Ptprc^−^Ilr7^+^*), c4_Cd8-Dadpl1-Tnaive (*Cd8^+^Ptprc^+^Ccr7^+^*), c5_Cd8-Mif-Tem (*Cd8^+^Ptprc^−^Prf1^+^*), and c8_NKT-Ccl5 (*Cd8^+^NKg7^+^*); five clusters for CD4^+^T cells: c0_Cd4-Fos-Tnaive (*Cd4^+^Ptprc^+^Ccr7^+^*), c1_Cd4-Lef1-Tcm (*Cd4^+^Ptprc^−^Ccr7^+^*), c3_Cd4-Fyn-Teffector (*Cd4^+^Ptprc^+^ Ccr7^−^*), c6_Cd4-Ifng-Th (*Cd4^+^Ptprc^−^Ifng^+^*), and c9_Cd4-Ctla4-Tex (*Cd4^+^Ctla4^+^*); and one cluster for innate lymphoid cells (ILCs): c7_ILC2-Rora (*Cd4^−^Cd8^−^Rora^+^*) (**Figure 4A, B**; Table S16).

**Figure 4.**
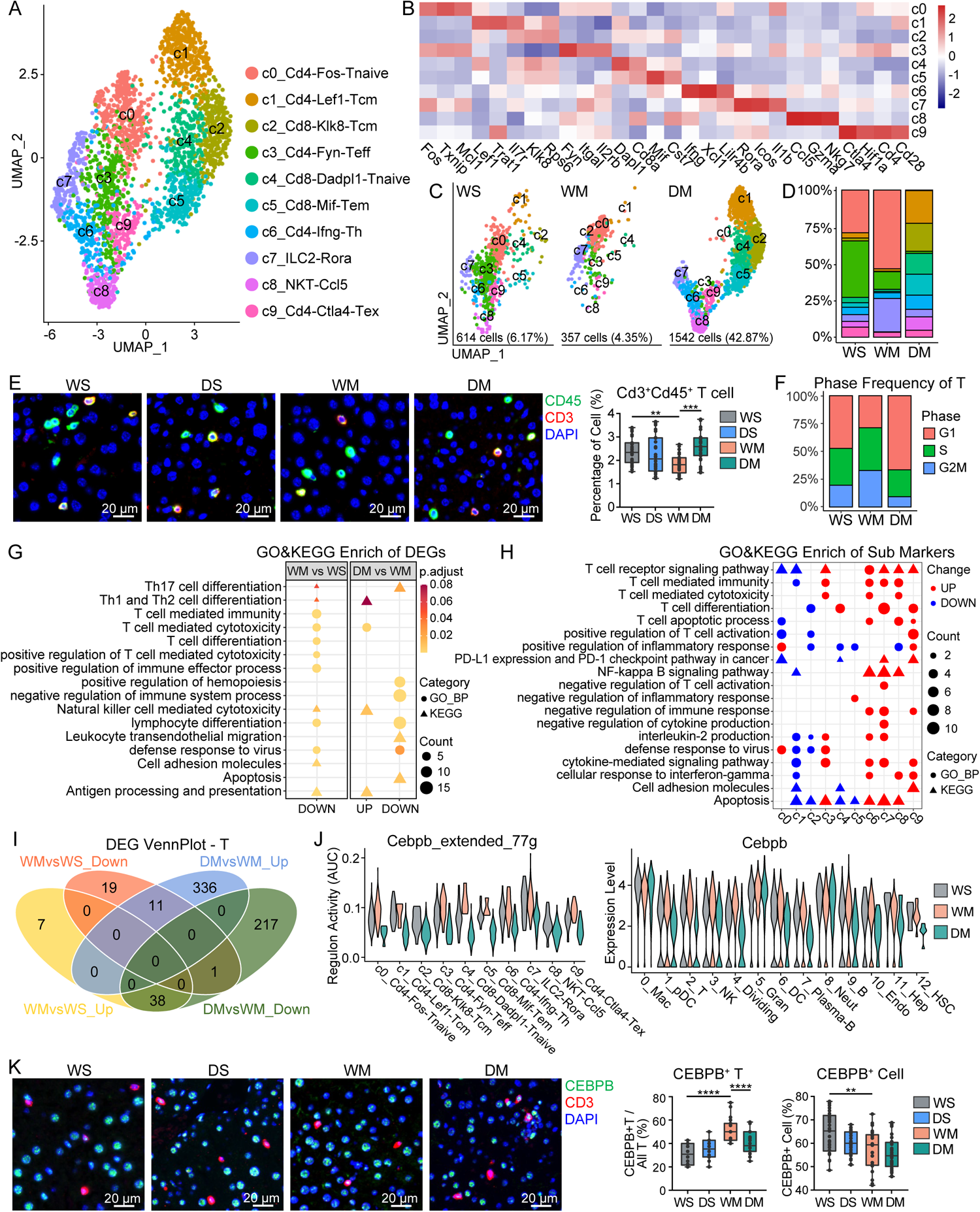
METH induces T-cell immunodeficiency. **A.** The 10 subclusters were identified from the T cells. **B.** The expressions of the marker genes for each T cell subcluster. **C. & D.** Group-wise cell populations and proportions of T cell. The legend is shared with (A). **E.** Representative double immunofluorescent of mice hepatic tissue (n=3) with DAPI nuclear counterstain (×40, bar = 20 μm) (left). Quantification of CD3+CD45+T cells (right). WS: WT + Saline; WM: WT + METH; DS: DRD1 KO + Saline; DM: DRD1 KO + METH. *: P < 0.05, ****: P < 0.0001. **F.** The percentage of cell cycle phases of T cells in 3 groups. **G.** GO and KEGG enrichment of the up-regulated and down-regulated genes of T cells. **H.** GO and KEGG enrichment of the marker genes of the T cell subclusters. **I.** The intersection of up-regulated and down-regulated genes of T cells. **J.** Regulon activity (AUC) of Cebpb and its targets (left) and expression level of Cebpb in all cell types (right). **K.** Representative double immunofluorescent of mice hepatic tissue (n=3) with DAPI nuclear counterstain (×40, bar = 20 μm) (left). Quantification of CEBPB+T cells and CEBPB+ cells (right), **: P < 0.01, ****: P < 0.0001.

The number of T cells from the DM group significantly increased compared with the WS and WM groups (**Figures 2B and 4C, D**), which was verified by immunofluorescence of the liver tissue (**Figure 4E**). We observed that the DM group had the most G_1_ cells and the least G_2_M cells among the three samples (**Figure 4F**), and the apoptosis pathway was down-regulated concurrently (**Figure 4G**). The proportion of CD8^+^T cells in the DM group increased significantly, but most of them were naive and memory cells except nature killer T cell (NKT). According to the enrichment results, DRD1 knockout eliminated some of the inhibition of METH on T cell differentiation and cytotoxicity. However, owing to the regulatory effect of DA on T cells [32, 33], there may be some changes that we did not observe.

The proportions of c0_Cd4-Fos-Tnaive and c7_ILC2-Rora were increased and that of c3_Cd4-Fyn-Teff was decreased in the WM group, whereas c0_Cd4-Fos-Tnaive and c3_Cd4-Fyn-Teff were almost absent in the DM group, with significant increases in subclusters c1_Cd4-Lef1-Tcm, c2_Cd8-Klk8-Tcm, c4_Cd8-Dadpl1-Tnaive, and c5_Cd8-Mif-Tem, which were non-active naive or memory cells. Enrichment and gene set variation analysis (GSVA) of the marker genes for each subcluster showed that the c0_Cd4-Fos-Tnaive subcluster is the naive cells, and c7_ILC2-Rora has the function of inhibiting T cell activation and immune response (**Figure 4H**; Figure S5A; Tables S17, 18). These findings indicate that METH may inhibit T cell activation and differentiation. The high expression of *Icos* in ILC2 has been reported to induce an immunosuppressive microenvironment [34]. The proportions of subclusters c0_Cd4-Fos-Tnaive and c7_ILC2-Rora decreased after DRD1 knockout. Subcluster c8_NKT-Ccl5 NKT cells have strong immune activity, which almost disappeared in the WM group but was present in the DM group. Subcluster c9_Cd4-Ctla4-Tex expresses high levels of exhaustion marker *Ctla4* and has some fragile regulatory T cell (Treg) characteristics (high expression of *Hif1a* and loss of suppressive function) [35] (Figure S5B), but no increase in the proportion of this cell subcluster was detected in the WM group.

Compared with the WS group, T cells in the WM group exhibited down-regulation of differentiation, cytotoxicity, defense response to virus, and antigen processing and presentation pathways. In the DM group, the pathways of cytotoxicity and antigen processing and presentation were up-regulated, but the defense response to virus pathway was still down-regulated (Figure 4G; Figure S5C; Tables S19, 20). Genes regulated by METH, and reversely regulated by DRD1 deletion were also found in T cells by VDA (**Figure 4I**; Table 1), including transcription factor *Cebpb* and effector (secreted protein) *Ccl5*. An interaction network of the T cell genes regulated by METH via DRD1, based on the STRING Database, was generated (Figure S5D and Table S21). The transcription factor analysis of T cells showed a higher expression level of *Cebpb* and its regulatory network in the WM group (mainly consisting of c0_Cd4-Fos-Tnaive and c7_Ilc2-Rora subclusters), with a lower expression level in the DM group (**Figure 4J**; Table S22). *Cebpb*, as an inhibitor of T cell proliferation, is involved in many immunological processes in other immune cells [36–38]. The increase of CEBPB^+^T and decrease of CEBPB in the WM group compared with the WS group may be related to the reduction of T cells and functional suppression of immune cells. The expression changes of *Cebpb* in T cells of the liver were verified by tissue immunofluorescence **(Figure 4K)**.

### METH causes DRD1-related immunosuppression of NK and B cells

The changes in proportion and cell cycle scores of 2053 NK cells in the three groups **(Figure 5A, B)** were similar to those of T cells. Compared with the WS group, NK cells in the WM group showed down-regulation of cytotoxicity, response to virus, and antigen processing and presentation pathways. In the DM group, these pathways were up-regulated, indicating that DRD1 knockout prevented the METH effect (**Figure 5C**; Tables S23, 24). In contrast, the apoptosis-associated pathways were up-regulated in the WM group and down-regulated in the DM group.

**Figure 5.**
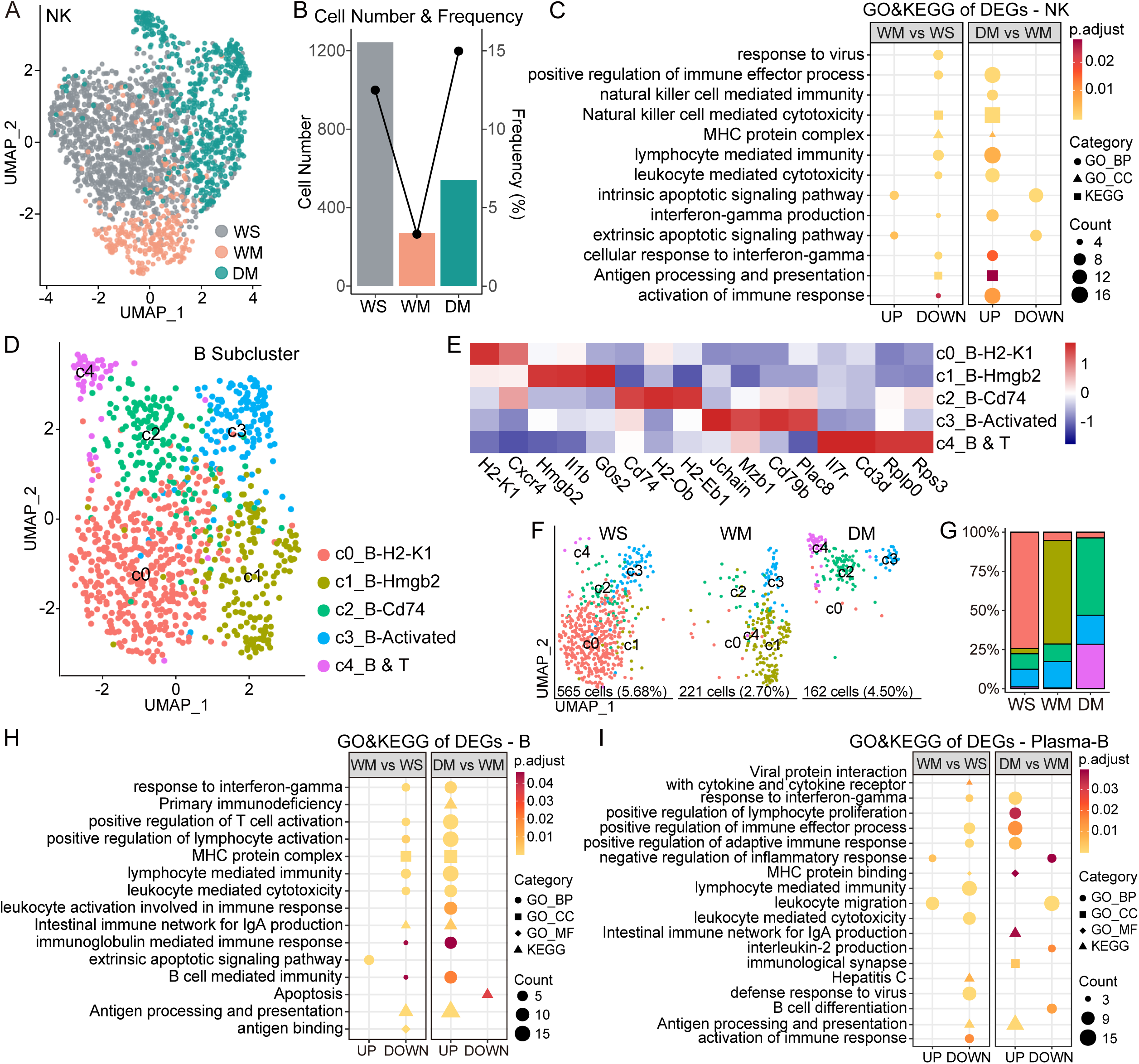
METH causes NK and B cells to exhibit DRD1-related immunosuppression. **A.** UMAP visualization of NK cells from 3 groups. **B.** Cell number and proportion of NK cells from 3 groups. **C.** GO and KEGG enrichment analysis of the up-regulated and down-regulated genes in NK cells of WM vs. WS group and DM vs. WM group. **D.** UMAP visualization of B cells showing a five-subcluster distribution. **E.** The expression of marker genes of B cell subclusters. **F. & G.** Group-wise cell populations and proportions of B cell. The legend is shared with (E). **H.** GO and KEGG enrichment of the up-regulated and down-regulated genes in B cells of WM vs. WS group and DM vs. WM group. **I.** GO and KEGG enrichment of the up-regulated and down-regulated genes in Plasma-B of WM vs. WS group and DM vs. WM group.

A total of 948 B cells were further subdivided into five subclusters (**Figure 5D**; Table S25). As a subcluster of naive B cells, c0_B-H2-K1 is functionally inactive and highly expresses *Cxcr4*. Subcluster c1_B-Hmgb2 displayed down-regulation of multiple immune functions and was present in an increased proportion in the WM group. As activated B cells with high expression of *Jchain*, subcluster c3_B-Activated showed up-regulation of B cell activation, differentiation, lymphocyte proliferation, activation of immune response, and response to IFN-gamma pathways. Subcluster c4_B&T, which highly expressed *Cd3d*, showed up-regulation of the response to interleukin-4 and immunological synapse pathways and was a heterozygous cell of activated B cells and T cells (**Figure 5E–G**, Figure S6, Table S26).

The proportion of B cells decreased in the WM group and increased in the DM group, with no significant difference in plasma B cells among the three groups (Figures 2B and 5F). Compared with the WS group, B cells in the WM group showed down-regulation of the pathways of antigen processing and presentation, B cell-mediated immunity, and response to IFN-gamma. Similar to T cells, B cells in the DM group showed recovery of some pathways that were down-regulated in the WM group, and a down-regulated apoptosis pathway (Figure 5G; Tables S27, 28). For plasma B cells, the changes in the WM group were similar to B cells. However, in the DM group, plasmacytes showed distinct down-regulation of the B cell differentiation pathway and up-regulation of the positive regulation of lymphocyte proliferation and immunological synapse pathways (**Figure 5H, I**; Tables S29, 30).

### METH inhibits cell crosstalk between macrophages and T cells

CellChat analysis was performed to evaluate the probability of immune cell–cell communication by integrating gene expression with prior knowledge of the interactions between signaling ligands, receptors, and their cofactors. Since we observed that the functions of macrophages and Kupffer cells appeared to be different in enrichment analysis, we analyze these cells separately here.

In the analysis of differential interaction strength, we found that—except for the increase in macrophages—METH resulted in decreased crosstalk of Kupffer, DCs, and all lymphocytes with T cells, and decreased crosstalk of Kupffer, pDC, T and NK cells with NK cells. However, DRD1 knockout led to the recovery of communication between T cells and other cells, but that of NK cells with Mac, T, and NK cells further decreased, indicating that METH regulated Mac, Kupffer, T, and NK cells differently **(Figure 6A)**.

**Figure 6.**
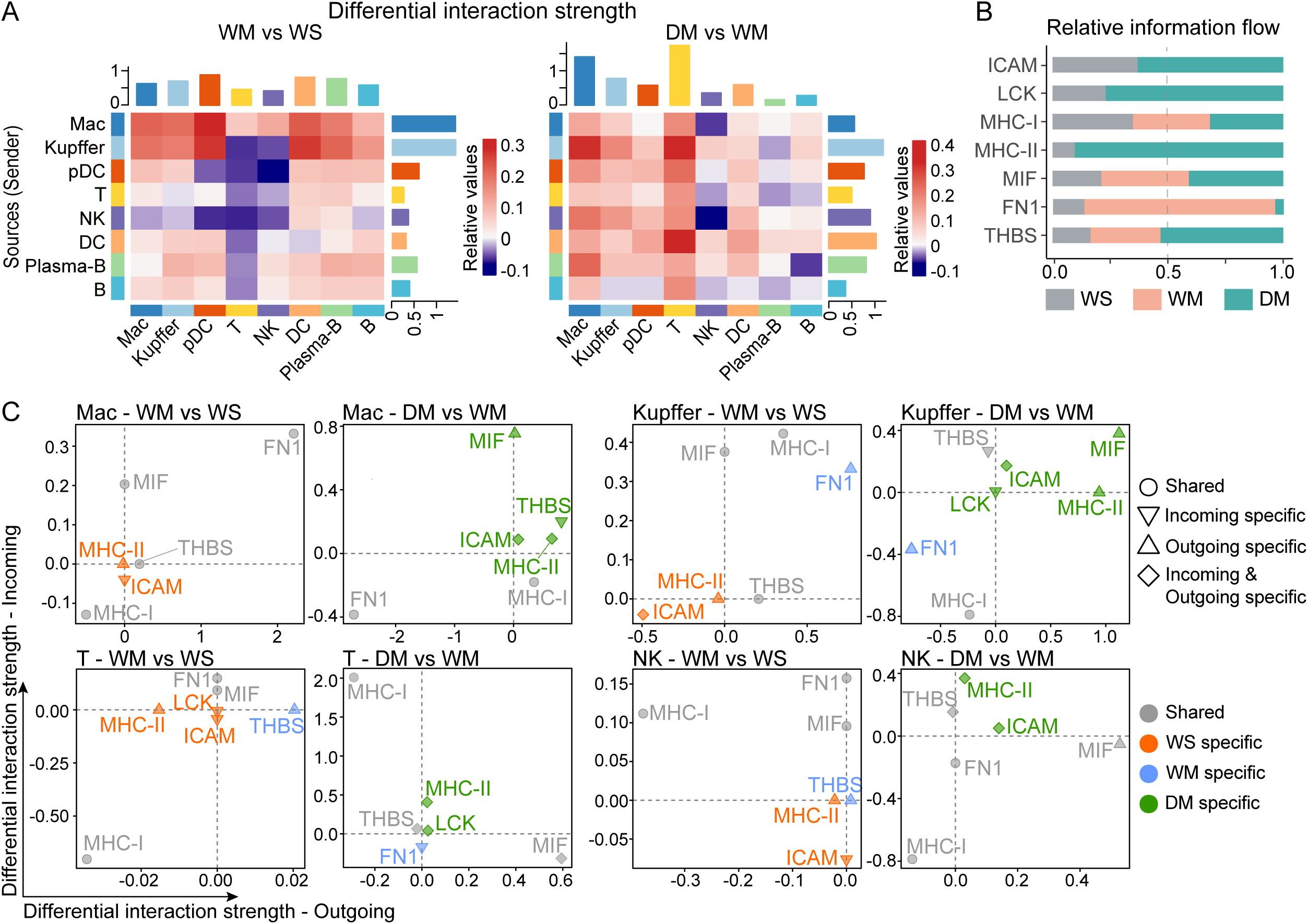
METH induces impaired cross-talk between immune cells. **A.** The differential interaction strength of 3 groups. Red (or blue) represents increased (or decreased) signaling in group WM compared to group WS (group DM compared to group WM). The top barplot represents the sum of the column of values displayed in the heatmap (incoming signaling). The right barplot represents the sum of the row of values (outgoing signaling). **B.** Relative information flow and ligand and receptor of interested pathways in 3 groups, which is defined by the sum of communication probability among all pairs of cell groups in the inferred network. **C.** Visualizing differential outgoing and incoming signaling changes in macrophage, Kupffer, T, and NK cells from WM to WS and DM to WM. Shapes of dots: circle, both groups have the outgoing and incoming signaling of the pathway; inverted triangle, only incoming signaling of the pathway was detected in one group; triangle, only outgoing signaling of the pathway was detected in one group; diamond, both outgoing and incoming signaling of the pathway were detected in one group, or one signaling of the pathway were detected in one group and another signaling in another group. Color of point: grey, shared pathway (detected in two groups); red, WS-specific pathway (lost in WM); blue, WM-specific pathway (lost in WM); green, DM-specific pathway (lost in WM).

In the analysis of the overall information flow of the pathway, the WM group lost ICAM (intercellular adhesion molecule), LCK (tyrosine-protein kinase Lck), and MHC II (main pathway triggering CD4 T cell receptor (TCR)) but exhibited a significant increase in the FN1 (fibronectin 1) pathway **(Figure 6B**; **Table 2)**. Further analysis of the pathways of the main cells that had changed **(Figure 6C)**, showed that: 1) The MHC II pathway disappeared in the WM group, and the MHC I signal mostly weakened, which was consistent with the down-regulation of the antigen processing and presentation pathway in most of the immune cells in this group; 2) In the WM group, the signal of the FN1 pathway in macrophages and Kupffer cells increased significantly, which may indicate an increase of immune infiltration or anti-inflammatory M2 [39]; 3) The ICAM signal disappeared in all cells in the WM group, which may lead to the weakening of the inhibition of T cell pro-inflammatory factors [40]; 4) The LCK signal disappears in T cells in the WM group, which may affect the initial triggering of TCR [41]; 5) The MIF (macrophage migration inhibition factor) signal increased in macrophages and Kupffer cells in the DM group. MIF is a pleiotropic cytokine with chemokine-like functions, which may be related to the decrease of macrophages and the increase of the proportion of B cells in the DM group [42, 43]. The details of the cell-to-cell communication analysis are listed in Table S31.

**Table 2.**
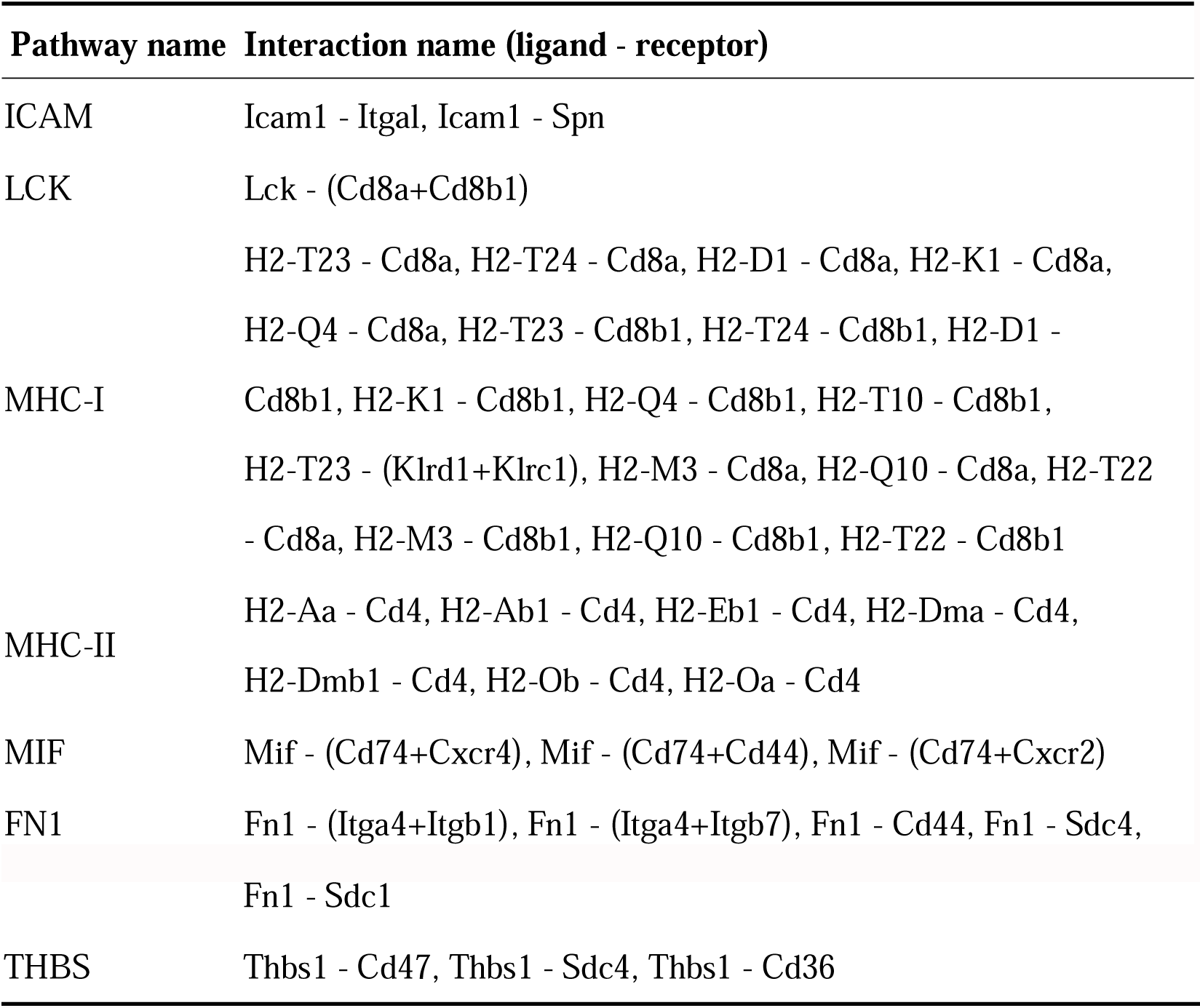
Ligand and receptor in the pathways.

## Discussion

In this study, by using scRNA sequencing, we described the changes of different liver-resident immune cells in response to METH and the possible role of DRD1 in this process. This dataset can provide new clues to therapies for METH-induced immune impairment.

Macrophages, as crucial elements of the innate immune system, can respond rapidly to various stimuli, such as infections with bacteria, viruses, or fungi, and other injuries. In this study, we found that chronic exposure to METH shifted the hepatic c0_Mac-Ifitm3 (named as Ifitm3^+^Mac here) and c2_Mac-Ccl5 (named as Ccl5^+^Mac here) into c1_Mac-Cd14 (named as Cd14^+^Mac here), with down-regulation of multiple immune functional pathways. This shift and down-regulation led to macrophage immune suppression in the WM group, supporting that METH exposure leads to suppression of hepatic immune functions. Among many decreased immune factors, the decrease in *Ifitm3* and *Ccl5* and increase in *Cd14* (also named monocyte differentiation antigen) in macrophages (Figure 3G; Supplementary Figure 4B) play a key role in the suppression of innate immunity by METH. *Ifitm3* can restrict various viral infections and is closely related to the antiviral ability of cells [44, 45]. CCL5 can facilitate M1 polarization and impede M2 polarization and is associated with liver injury [46]. CD14, which is predominantly found in monocytes and macrophages, is a marker of myeloid differentiation and inactivation. The surface expression of CD14 on monocytes is reduced after cell activation [47]. These results indicate that chronic exposure to METH impaired the antiviral or immune capacity of macrophages. In addition, we found that the proportion of macrophages in the DM group almost disappeared, which was consistent with the immunofluorescence results (Figure 3G). Apart from the reason of G_1_ cell cycle arrest (Supplementary Figure 4A), we believe that this loss of macrophages is the result of DRD1 deletion rather than an effect of METH. The loss of macrophages and its mechanism need to be confirmed and investigated further.

DA binds to D1-like receptors expressed by mouse Tregs and can inhibit the inhibitory activity of Tregs on effector T cells [48]. The concentration of DA can also regulate the proliferation of T cells. At a physiological concentration range of µM–nM (10^−9^–10^−6^ M), DA usually suppresses activated effector T cells, leading to inhibition of their proliferation, cytokine secretion, and other processes and responses. However, at a much higher concentration of approximately 0.1–1 mM (10^−4^–10^−3^ M), DA can be toxic and kill the T cells [32]. The administration of METH can result in a marked increase in extracellular DA [19]. We speculate that the changes in T cell numbers in the WM and DM groups might be related to an increase in DA concentration due to METH and DRD1 knockout, respectively. In addition, the proportion of macrophages in the DS and DM groups decreased significantly, indicating that DRD1 is required for hepatic macrophage homeostasis, but further research is needed to confirm this theory.

In our study, chronic exposure to METH inhibited the functions or proportion of most immune cells in the liver, leading to the down-regulation of some immune-related pathways. Some of these pathways were restored in the DRD1 knockout group (DM), proving the involvement of DRD1 in the suppressive effects of METH. Among these pathways, the antigen processing and presentation pathway was the most significant (Figures 3G, 4E, and 5C, G), especially in the communication between T cells and macrophages or other antigen-presenting cells through MHCII, LCK, and ICAM, and in the communication between T and NK cells through LCK. These findings demonstrated that the structure and the function of DRD1 was associated with MHCII, LCK, and ICAM signaling pathways (Figure 6C, Supplementary Table 31). We speculate that the main mode of METH affecting liver immunity may be via DRD1–MHCII/LCK. The mechanism of METH on immune-related pathways dependent or independent of DRD1 needs to be explored further.

Lymphocytes, especially T cells, are the main cells involved in adaptive immunity against viruses or other infections. METH is likely to cause disorders of other immune organs besides the liver [5], as shown for opioid addiction in previous studies. Opioid-associated blood exhibited an abnormal distribution of immune cells, characterized by significant expansion of fragile Tregs, and showed enhanced Treg-derived IFN-gamma expression [49]. In our study, there was a similar *Foxp3^+^* subcluster (c9_Cd4-Ctla4-Tex) that lost its inhibitory function (Figure 4B). However, this subcluster had high expression of *Hif1a* but did not express IFN-gamma (Supplementary Figure 5B). Furthermore, our results showed that METH exposure decreased the ratio of CD8^+^T cells in the liver and inhibited a series of functions such as cell differentiation, proliferation, and cytotoxicity (Figure 4D, G). This discrepancy between our findings and those of studies with opioids may be related to the different effects and mechanisms of opioids and METH. Opioids are central nervous system inhibitors that act through opioid receptors, whereas METH is a central nerve stimulant that increases extracellular DA levels through interactions with dopamine transporter [19]. The mechanisms of addiction of the two drugs are similar (related to DA and reward circuits), but the mechanism of their effects on the immune system may be different.

In our study, we found that chronic exposure to METH shifted the hepatic immune active c4_Cd4-Fyn-Teff (also named Fyn^+^CD4^+^Teff), CD8T, and NKT (c8-NKT) to c0_CD4-Fos-Tnaive (also named Fos^+^CD4^+^T) and c7_ILC2-Rora (also named Rora^+^ILC2), and inhibited the function of B cells and NK cells, leading to lymphocyte immune suppression. Our data revealed that METH may regulate many effectors via DRD1, in which GZMA and GZMB (Granzyme A and B) play a key role (Supplementary Figure 7). Cytotoxic T lymphocytes and NK cells use perforin to deliver serine protease granzymes into target cells to kill them [50]. The decrease of granzyme in the WM group in our study confirmed the decrease in cellular immune capacity.

This study has some limitations. First, hepatic NPCs were isolated predominantly, which restricts understanding of the crosstalk between liver parenchymal cells and immune cells. Second, bulk RNA-Seq revealed some effects of DRD1 knockout on liver immune function. However, the DS group was not included in scRNA-Seq experiments as we focused on METH, thus the effects of DRD1 knockout alone on hepatic immunity at the single-cell level are not known. Third, the alterations of some cell proportions, such as macrophages and T and B cells, in scRNA-Seq were not completely consistent with those observed in other experiments, such as immunofluorescence, flow cytometry, or ImmuCellAI estimation in bulk RNA-Seq. This discrepancy may be due to differences between mRNA and protein or differences in the technologies themselves. Although scRNA-Seq is a highly effective method for cellular-level analysis and visualization of the cellular landscape, the interpretation of its results is not yet consistent and precise owing to technical constraints and the varying algorithms used for data analysis across different studies.

In conclusion, we examined the effect of chronic exposure to METH on hepatic immune cells and investigated whether DRD1 was involved by using a METH exposure mouse model and DRD1 knockout mice with scRNA-Seq. Chronic exposure to METH shifted the immune cells from Ifitm3^+^Mac and Ccl5^+^Mac to Cd14^+^Mac, and from Fyn^+^CD4^+^Teff, CD8T, and NKT to Fos^+^CD4^+^T and Rora^+^ILC2, coupled with suppression of multiple immune functional pathways, and these effects were partially prevented by DRD1 deletion. Based on our findings, we consider that METH may suppress hepatic macrophages mainly through DA–DRD1/MHCII-IRF7-IFN/CCL5/others, and T cells by DA–DRD1/MHCII-CEBPB-CCL5/other effectors.

## Materials and methods

### METH chronically exposed mouse model

The experimental animals were provided by Professor Ming Xu (University of Chicago). Genotypes of the DRD1 knockout and their matched wild-type mice were determined by PCR using gene-specific primers as previously reported [51]. All mice were housed in a pathogen-free animal facility at Xi’an Jiaotong University, and the temperature and humidity of the room were controlled. Mice were group-housed in a 12-h light/dark cycle with free access to food and water.

The METH chronically exposed mouse model was modified based on a previous publication [52]. Briefly, METH was dissolved in sterile 0.9% physiological saline. Male 8-week-old wild-type (W set) and DRD1 knockout mice (D set) were randomly divided into four groups (two groups per set, *n* = 8) and received intraperitoneal injections of saline (groups WS and DS) or METH (groups WM and DM) for 2 weeks. During the initial 7 days, the mice received daily injections of 1 mg/kg of body weight, and for the subsequent 7 days, the dose was 2 mg/kg of body weight. Under deep anesthesia, mice were sacrificed 24 h after the last injection. The liver was dissected and stored by different methods according to subsequent experiments. METH (purity of 99.1%, identified by the National Institute for Food and Drug Control, Guangzhou, China) was obtained from the National Institute for the Control of Pharmaceutical and Biological Products (Beijing, China).

### Bulk RNA-Seq and data processing

The liver tissues were used for bulk RNA-Seq as described previously [53]. Briefly, total RNA was extracted from liver tissues and a cDNA library was constructed using a MGIEasy mRNA libraries kit V2 (*n* = 6 per group). We used limma (version 3.52.4), edgeR (version 3.38.4), and DESeq2 (version 1.36.0) to analyze differentially expressed genes (DEGs) [54–57], and took intersection as the final result. DEGs based on the criteria of *P* < 0.05 and log2(fold change) (log2FC) >1 were identified. DEGs enrichment analysis was performed using clusterProfiler (version 4.4.4) [58]. Immune infiltration analysis was performed by using ImmuCellAI-mouse (version 0.1.0) [59].

### Liver nonparenchymal cell isolation

Primary hepatic nonparenchymal cells (NPCs) were isolated using the digestion solution PSCeasy dispase (Cellapy, Beijing, China) according to the manufacturer’s instructions, as described previously [60]. Briefly, the liver was perfused with perfusion solution (calcium-free Hanks with 0.5 M EDTA) for 5 min. The liver was then minced and digested in PSCeasy solution at 37 □ for 20 min. The disassociated cells were filtered through a 40-μm cell strainer (Falcon, One Riverfront Plaza, NY, USA), then the immune cells and liver cells were separated by centrifugation at 50 *g* and 4 □ for 5 min. Erythrocytes were lysed with red blood cell lysis buffer (Beyotime, Shanghai, China) for 2 min and washed twice. The liver NPCs were filtered to remove dead cells before they were captured using a 10x Chromium Controller (10x Genomics).

### Flow cytometry

The liver cells were collected by gently mashing liver tissue through a 45-μm cell strainer, without liver perfusion. And then the cells and cell debris were separated by Percoll. Erythrocytes were lysed as described above. Flow cytometry was used for cell type analysis of NPCs as described previously [61]. Briefly, the isolated cells were centrifuged at 4 L, 350 *g* for 5 min. The cells were then washed and re-suspended in cold phosphate-buffer saline, followed by incubation with anti-CD16/CD32 antibody to block Fc receptors. Fluorochrome-conjugated antibodies against CD11b (APC), F4/80 (Brilliant Violet 421), CD8a (APC/Cyanine7), CD4 (PE), and CD3 (Alexa Fluor 488) were used to distinguish cell types. All antibodies were from Biolegend (San Diego, CA, USA). Macrophages were gated as CD11b^+^F4/80^+^; Cd4^+^ T cells were gated as CD3^+^CD4^+^CD8^−^; and Cd8^+^ T cells were gated as CD3^+^CD8^+^CD4^−^. Samples were analyzed using BD FACSAria™ Fusion Flow Cytometer. Data were analyzed using FlowJo software (Flowjo.com).

### Single-cell RNA-Seq and data processing

We used Cell Ranger version 2.0.1 (10x Genomics) to pool and process the raw RNA sequencing data. All reads were aligned to the mouse transcriptome reference genome (Mus_musculus_GRCm38.p5) through the Cell Ranger count pipeline. After data aggregation, we performed all filtering, normalization, and scaling of data using Seurat (version 4.1.0) [62, 63]. Cells with fewer than 200 and more than 3000 detected genes, as well as cells with fewer than 600 unique molecular identifiers and more than 5% mitochondrial counts were filtered out. Genes that were detected in fewer than 10 cells were removed. Gene counts for each cell were normalized by total expression, multiplied by a scale factor of 10000, and transformed into a log scale. Principal component analysis based on the highly variable genes detected (dispersion of 2) was performed for dimension reduction, and the top 35 principal components were selected. The clusters were visualized using Uniform Manifold Approximation and Projection (UMAP). The top expressing genes for each cluster and the DEGs between three groups (WS, WM and DM) were defined by the Seurat FindMarkers function with the Wilcoxon Rank Sum test. Cell-type-specific gene signatures were determined from the overlap of more highly expressed and canonical gene markers. DEGs based on the criteria of adjusted *P* values *P*.adj) < 0.05 and │log2FoldChange (log2FC)│ > 0.5 were identified. Enrichment analyses of the DEGs, including Gene Ontology (GO), Kyoto Encyclopedia of Genes and Genomes (KEGG), and Gene set enrichment analysis (GSEA), were performed using clusterProfiler (version 4.4.4) [58]. Intercellular crosstalk was inferenced and analyzed by CellChat (version 1.6.1) [64], which predicts major signaling inputs and outputs for cells through integrating gene expression with prior knowledge of the interactions between signaling ligands, receptors, and their cofactors. Transcription factors and their target genes were reconstructed and termed “regulons” by SCRNIC (version 1.2.0) [65].

### Immunofluorescence

Liver tissues from WS, DS, WM, and DM groups (*n* = 3 per group) were fixed in situ with 4% paraformaldehyde and embedded in paraffin. The methods for staining, scanning, and observation of sections are as described previously [60]. Primary antibodies used in the immunofluorescence analysis were CD3, CD45, F4/80, CD14, and IFITM3 (Servicebio, Wuhan, China). All primary antibodies were rabbit anti-mouse and were detected with Alexa Fluor® 488-conjugated or Cy3-conjugated goat anti-rabbit IgG (Servicebio). Sections were scanned with Pannoramic DESK (3D HISTECH, Budapest, Hungary). The browser software Caseviewer (version 2.4) was used to acquire images (×40). The proportion of cells is the ratio of the number of positive cells to the number of all nuclei in the image. GraphPad Prism (version 9.0) was used to calculate the statistical significance of differences in the immunofluorescence data using analysis of variance (ANOVA) at a 95% confidence level. *P* < 0.05 indicated statistical significance.

## Authors’ contributions

SBL and XML conceived the project. SBL and XML designed and supervised research. JTZ was responsible for the experiments, data collection and data analysis with help from XHL, CC, and JNF. JTZ performed bioinformatic analyses with YGX’s supervision. XML, YGX and JTZ finished the manuscript. SBL and JY provided support in writing, review and editing the manuscript. All authors read and approved the final manuscript.

## Data availability

The accession number for the sequencing raw data and processed data in this paper is GEO: GSE247843 (private accesses). The R code used to perform analysis are available at GitHub https://github.com/XiaomingLi-xjtu/METH-DRD1-Lv to ensure the replicability and reproducibility of these results.

## Ethics statement

The animal study was reviewed and approved by Xi’an Jiaotong university health science center.

## Acknowledgements

We thank Professor Ming Xu (University of Chicago) for the kind donation of DRD1 knockout mice. We thank postgraduate Wencai Wu (School of Life Science and Technology, Xi’an Jiaotong University) for his assistance in bioinformatics analysis.

## Supplementary material

### Legends for Supplementary

**Supplementary Figure 1 METH changed immune cells and immune-associated pathways in mice liver. A.** RNA-Seq revealed changes in the immune-associated pathways in mice livers. WS: WT + Saline, n=6; WM: WT + METH, n=6; DS: DRD1 KO + Saline, n=6; DM: DRD1 KO + METH, n=6. **B.** Immune cell abundance scores estimated by ImmuCellAI-mouse based on RNA-Seq data showed that the frequency of many immune cells may be changed after METH treatment. **C.** Frequency of different cell types in 4 groups by flow cytometry. Group information was the same as A. *: *P* < 0.05, ***: *P* < 0.001.

**Supplementary Figure 2 Dividing cell and its subclusters A.** The cell cycle phases for all clusters. **B. & C.** Group-wise cell populations and proportions of 4_Dividing cell. **D.** The expression profiles of Mki67 in all cells and marker genes of the 4_Dividing cell population.

**Supplementary Figure 3 Genes changed in the representative functional pathways** Heatmap showing the relative gene expressions in the representative functional pathways.

**Supplementary Figure 4 DEGs of macrophages and Kupffer cell A.** Cell cycle phases of macrophages from 3 groups. **B.** Volcano plots showing fold change of gene expression (log2 scale) for down-regulated and up-regulated genes in Kupffer cells of WM vs. WS group and DM vs. WM group. Up-regulated genes (*P.adj* < 0.05 and log2FC > 0.5) are shown with red dots, down-regulated genes (*P.adj* < 0.05 and log2FC < −0.5) shown with blue dots, and insignificant genes (*P.adj* > 0.05 or log2FC < 0.5) shown with gray dots. **C.** Volcano plots showing fold change of gene expression (log2 scale) for down-regulated and up-regulated genes in Kupffer cells of WM vs. WS group and DM vs. WM group. Up-regulated genes (*P.adj* < 0.05 and log2FC > 0.5) are shown with red dots, down-regulated genes (*P.adj* < 0.05 and log2FC < −0.5) shown with blue dots, and insignificant genes (*P.adj* > 0.05 or log2FC < 0.5) shown with gray dots. **D.** GO and KEGG enrichment of the up-regulated and down-regulated genes of Kupffer cells in (C). **E.** Heatmap showing the relative gene expression levels in the representative immune pathways of macrophages from 3 groups. **F.** STRING networks of the gene sets regulated by METH through DRD1 in macrophages.

**Supplementary Figure 5 T subclusters function and DEGs A.** GSVA (gene set variation analysis) of T-cell subclusters. **B.** The expressions of the feature genes from the c9_Cd4-Ctla4-Tex subcluster. **C.** Volcano plots showing fold change of gene expression (log2 scale) for down-regulated and up-regulated genes in T cells of WM vs. WS group and DM vs. WM group. Up-regulated genes (*P.adj* < 0.05 and log2FC > 0.5) are shown with red dots, down-regulated genes (*P.adj* < 0.05 and log2FC < −0.5) shown with blue dots, and insignificant genes (*P.adj* > 0.05 or log2FC < 0.5) shown with gray dots. **D.** STRING networks of the gene sets regulated by METH through DRD1 in T cells.

**Supplementary Figure 6 Enrichment of the marker genes of B subclusters** GO and KEGG enrichment analysis of marker genes of five subclusters of B cells.

**Supplementary Figure 7 The expression levels of *Gzma* and *Gzmb*** The expression levels of *Gzma* and *Gzmb* in all cell types from group WS, WM and DM.

### Legends for Supplementary

**Supplementary Table 1. DEGs of RNAseq.** DEGs were analyzed by limma, edgeR, and DESeq2 respectively, and took intersection as the final result.

**Supplementary Table 2. Enrichment of DEGs of RNAseq.** GO and KEGG enrichment analysis of DEGs in Supplementary Table 1. Define genes with *P.adj* < 0.05 and log2FC > 1 as upregulated genes, and *P.adj* < 0.05 and log2FC < −1 as downregulated genes.

**Supplementary Table 3. Top50 markers of all cell type.** Top50 marker genes of all cell type, sort by log2FC.

**Supplementary Table 4. Cell number and frequency of all cell type from 3 group.** Cell number and frequency (in brackets) of all cell type from group WS, WM and DM.

**Supplementary Table 5. DEGs of scRNA – group.** Group-wise DEGs of scRNA.

**Supplementary Table 6. Enrichment of DEGs of scRNA – group.** GO and KEGG enrichment analysis of DEGs in Supplementary Table 5. Define genes with *P.adj* < 0.05 and log2FC > 0.5 as upregulated genes, and *P.adj* < 0.05 and log2FC < −0.5 as downregulated genes.

**Supplementary Table 7. GSEA enrichment of DEGs of scRNA – group.** GSEA enrichment analysis of all DEGs (*P.adj* < 0.05) in Supplementary Table 5.

**Supplementary Table 8. Top30 markers of 0_Mac subclusters.** Top30 marker genes of subclusters of 0_Mac, sort by log2FC.

**Supplementary Table 9. DEG of scRNA - Macrophage (except subcluster3).** Group-wise DEGs of subclusters c0, c1, c2, c4 and c5 of 0_Mac.

**Supplementary Table 10. Enrichment of DEGs - Macrophage (except subcluster3).** GO and KEGG enrichment analysis of DEGs in Supplementary Table 9. Define genes with *P.adj* < 0.05 and log2FC > 0.5 as upregulated genes, and *P.adj* < 0.05 and log2FC < −0.5 as downregulated genes.

**Supplementary Table 11. Enrichment of subcluster markers - Mac Subclusters.** GO and KEGG enrichment analysis of marker genes of six subclusters of 0_Mac.

**Supplementary Table 12. DEG of scRNA – Kupffer.** Group-wise DEGs of subcluster c3_Kupffer-Lgmn of 0_Mac.

**Supplementary Table 13. Enrichment of DEGs – Kupffer.** GO and KEGG enrichment analysis of DEGs in Supplementary Table 12. Define genes with *P.adj* < 0.05 and log2FC > 0.5 as upregulated genes, and *P.adj* < 0.05 and log2FC < −0.5 as downregulated genes.

**Supplementary Table 14. Intersection of DEGs – Macrophage.** Intersection of group-wise DEGs in Supplementary Table 9 and 12.

**Supplementary Table 15. Transcription factors analysis: regulon targets information – Macrophage.** Regulon targets information of transcription factors analysis of 0_Mac.

**Supplementary Table 16. Top30 markers of T subclusters.** Top30 marker genes of subclusters of 2_T, sort by log2FC.

**Supplementary Table 17. Enrichment of subcluster markers - T subclusters.** GO and KEGG enrichment analysis of marker genes of ten subclusters of 2_T.

**Supplementary Table 18. GSVA of T subclusters.** Results of GSVA (gene set variation analysis) of T-cell subclusters.

**Supplementary Table 19. DEGs of scRNA – T.** Group-wise DEGs of T cells.

**Supplementary Table 20. Enrichment of DEGs – T.** GO and KEGG enrichment analysis of DEGs in Supplementary Table 19. Define genes with *P.adj* < 0.05 and log2FC > 0.5 as upregulated genes, and *P.adj* < 0.05 and log2FC < −0.5 as downregulated genes.

**Supplementary Table 21. Intersection of DEGs – T.** Intersection of group-wise DEGs in Supplementary Table 19.

**Supplementary Table 22. Transcription factors analysis: regulon targets information – T.** Regulon targets information of transcription factors analysis of T cells.

**Supplementary Table 23. DEGs of scRNA – NK.** Group-wise DEGs of NK cells.

**Supplementary Table 24. Enrichment of DEGs – NK.** GO and KEGG enrichment analysis of DEGs in Supplementary Table 23. Define genes with *P.adj* < 0.05 and log2FC > 0.5 as upregulated genes, and *P.adj* < 0.05 and log2FC < −0.5 as downregulated genes.

**Supplementary Table 25. Top30 markers of B subclusters.** Top30 marker genes of subclusters of B cells, sort by log2FC.

**Supplementary Table 26. Enrichment of subcluster markers - B subclusters.** GO and KEGG enrichment analysis of marker genes of five subclusters of 9_B.

**Supplementary Table 27. DEGs of scRNA – B.** Group-wise DEGs of B cells.

**Supplementary Table 28. Enrichment of DEGs – B.** GO and KEGG enrichment analysis of DEGs in Supplementary Table 27. Define genes with *P.adj* < 0.05 and log2FC > 0.5 as upregulated genes, and *P.adj* < 0.05 and log2FC < −0.5 as downregulated genes.

**Supplementary Table 29. DEG of scRNA - Plasma-B.** Group-wise DEGs of Plasma-B.

**Supplementary Table 30. Enrichment of DEGs - Plasma-B.** GO and KEGG enrichment analysis of DEGs in Supplementary Table 29. Define genes with *P.adj* < 0.05 and log2FC > 0.5 as upregulated genes, and *P.adj* < 0.05 and log2FC < −0.5 as downregulated genes.

**Supplementary Table 31. Cell Chat: All inferred cell-cell communications.** The full list of predicted ligand - receptor pairs.

