## Supplementary figures and images for "Single-cell RNA Sequencing Reveals Methamphetamine Inhibits the Liver Immune Response with Involvement of the Dopamine D1 Receptor"

### Figure S1

Supplementary Figure 1

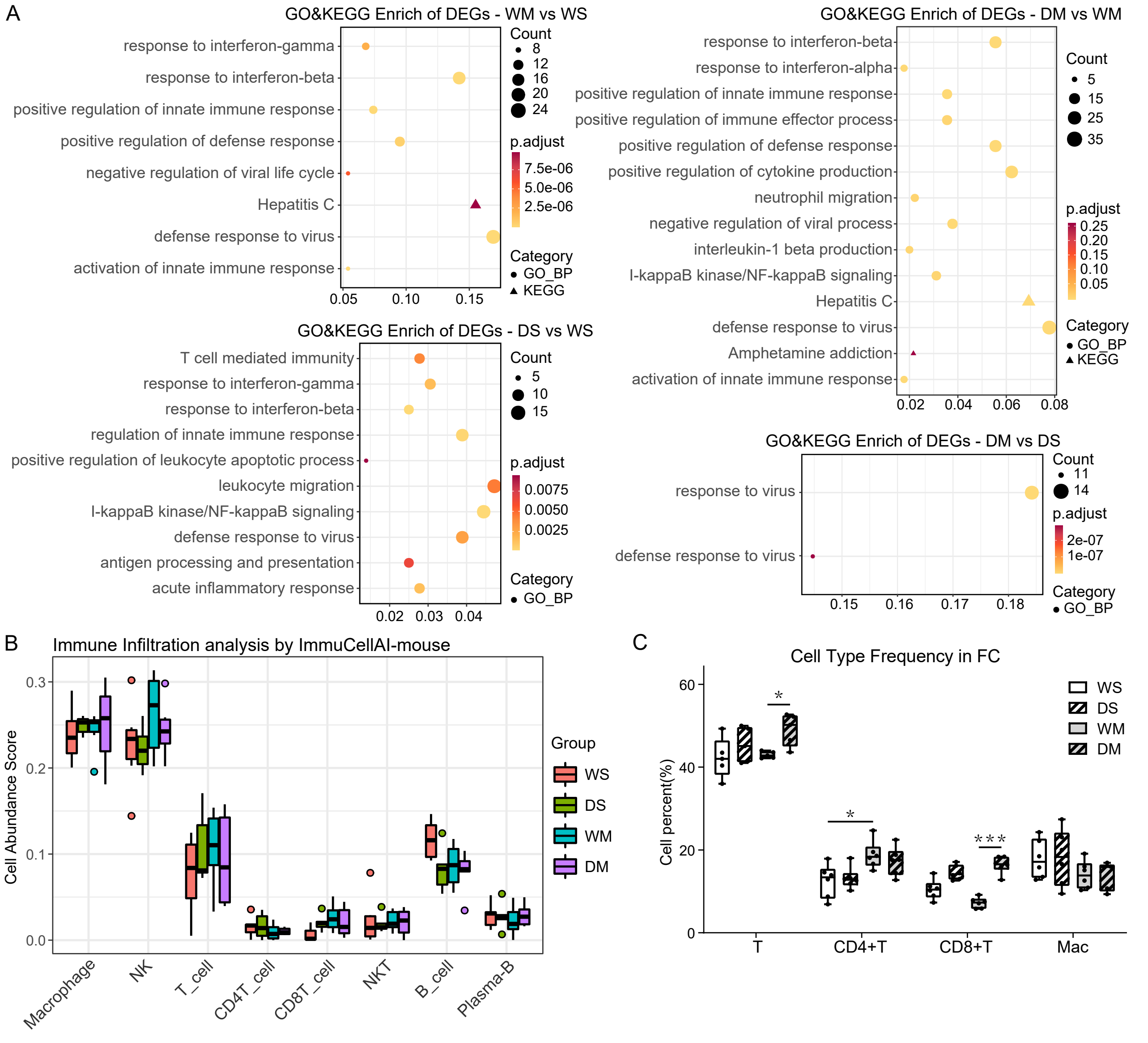

### Figure S2

Supplementary Figure 2

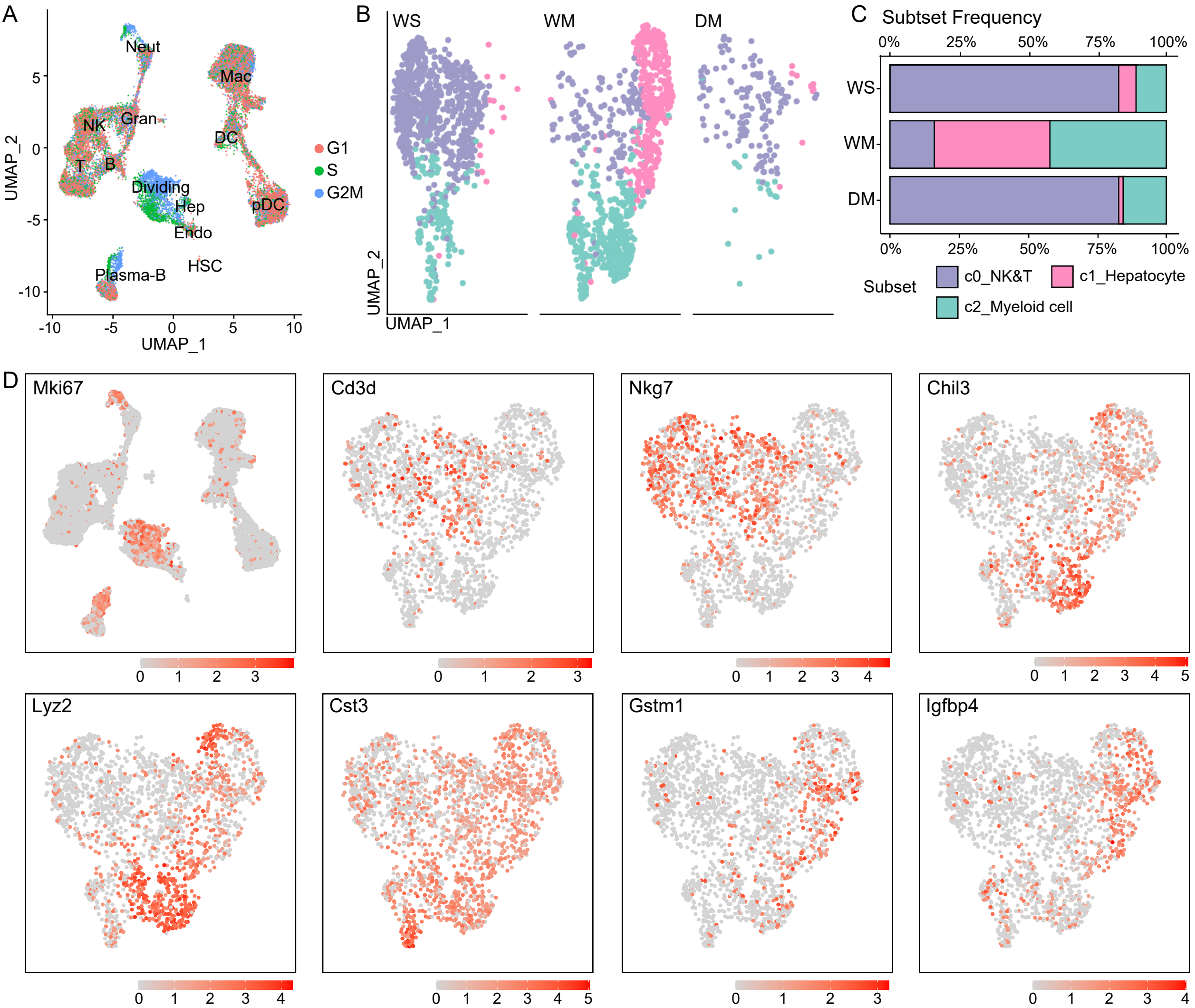

### Figure S3

Supplementary Figure 3

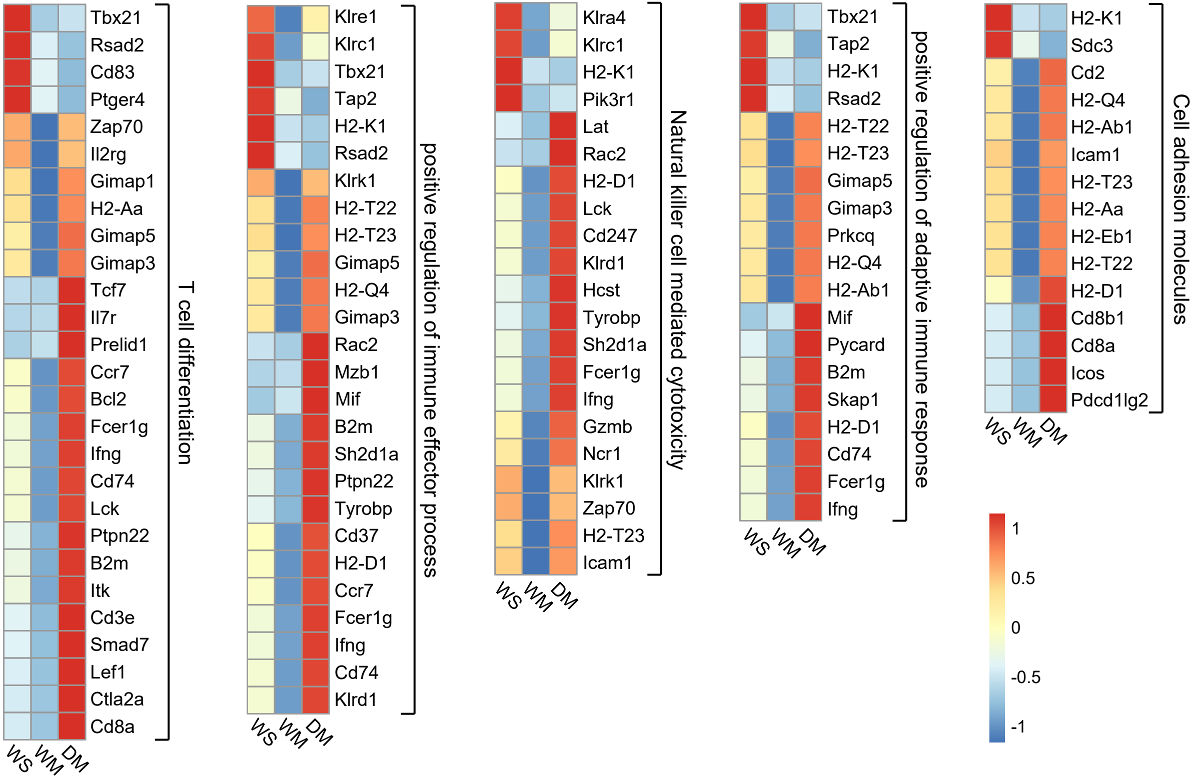

### Figure S4

Supplementary Figure 4

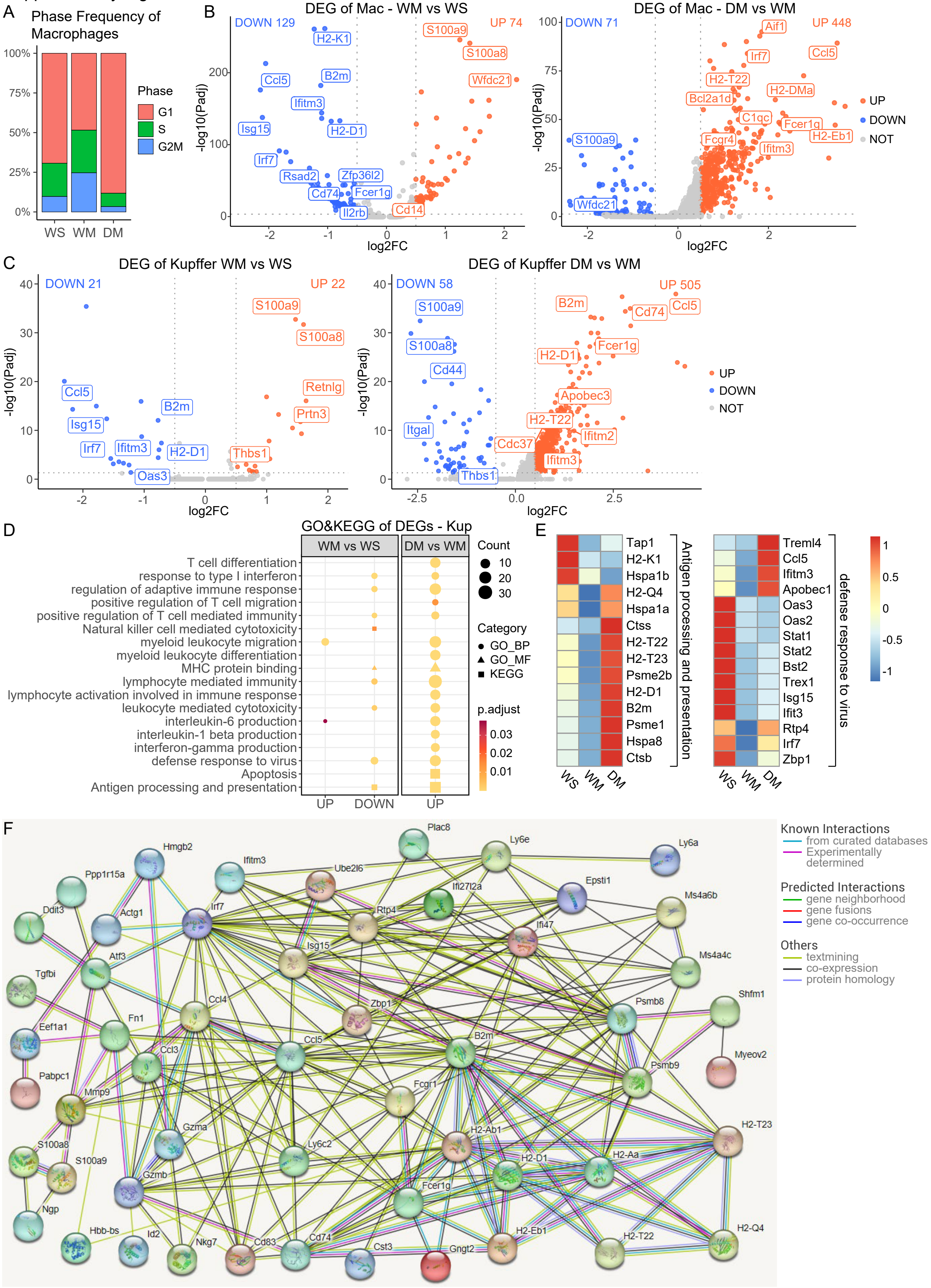

### Figure S5

Supplementary Figure 5

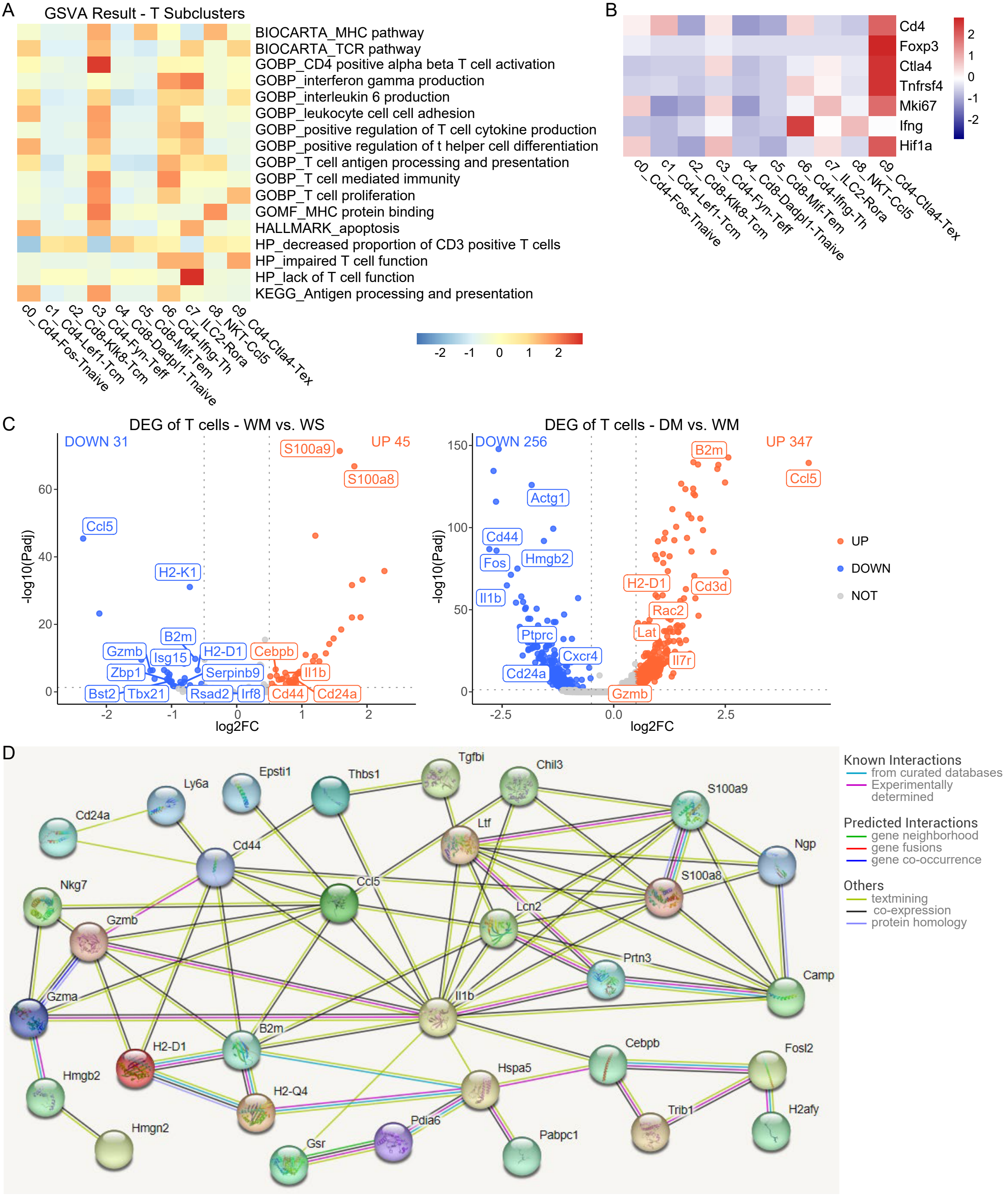

### Figure S6

Supplementary Figure 6

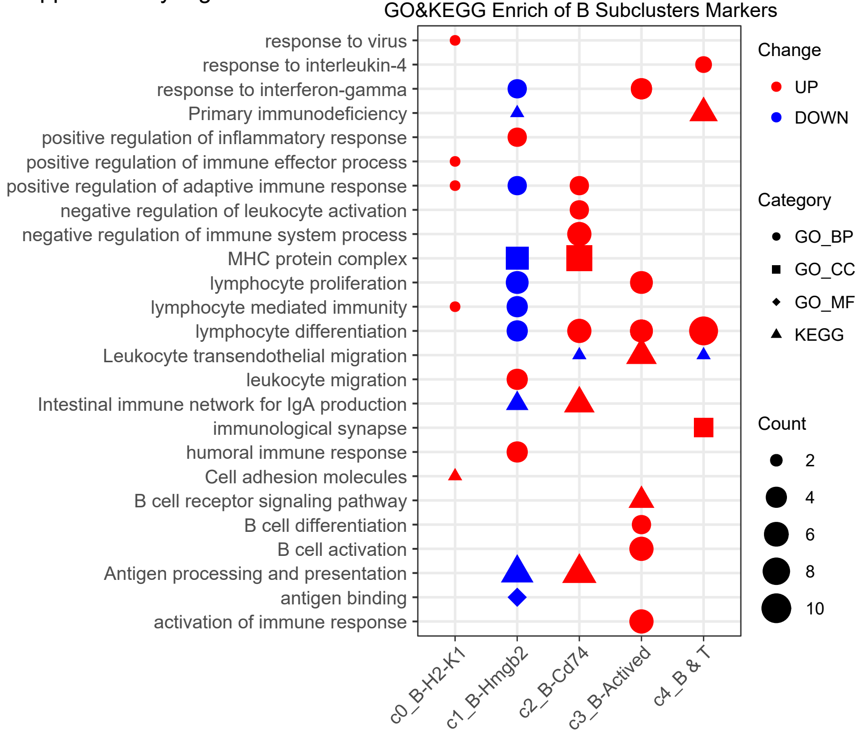

### Figure S7

Supplementary Figure 7

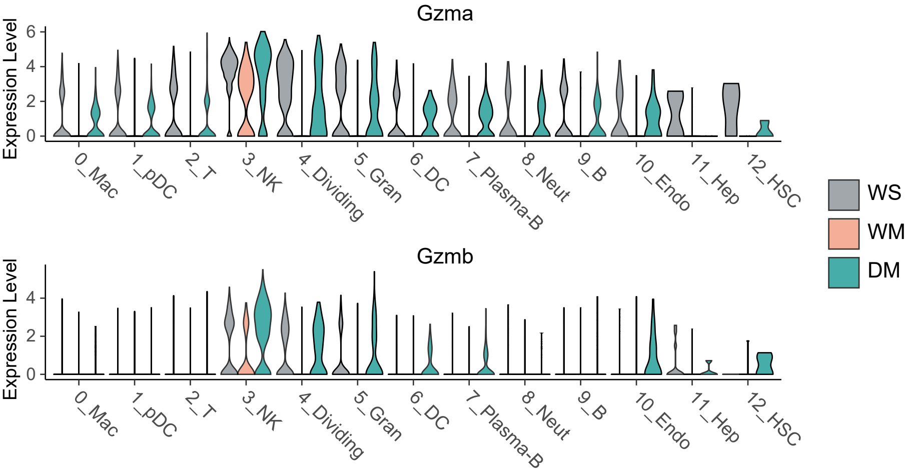
