## Supplementary material for "Single-cell RNA Sequencing Reveals Methamphetamine Inhibits the Liver Immune Response with Involvement of the Dopamine D1 Receptor": Legend of Supplementary Figure and Tables

### 1 This file includes:

Supplementary Figures S1–S7.

Legends for Supplementary Tables S1–S31.

### 2 Supplementary Figures and Tables

#### 2.1 Supplementary Figures

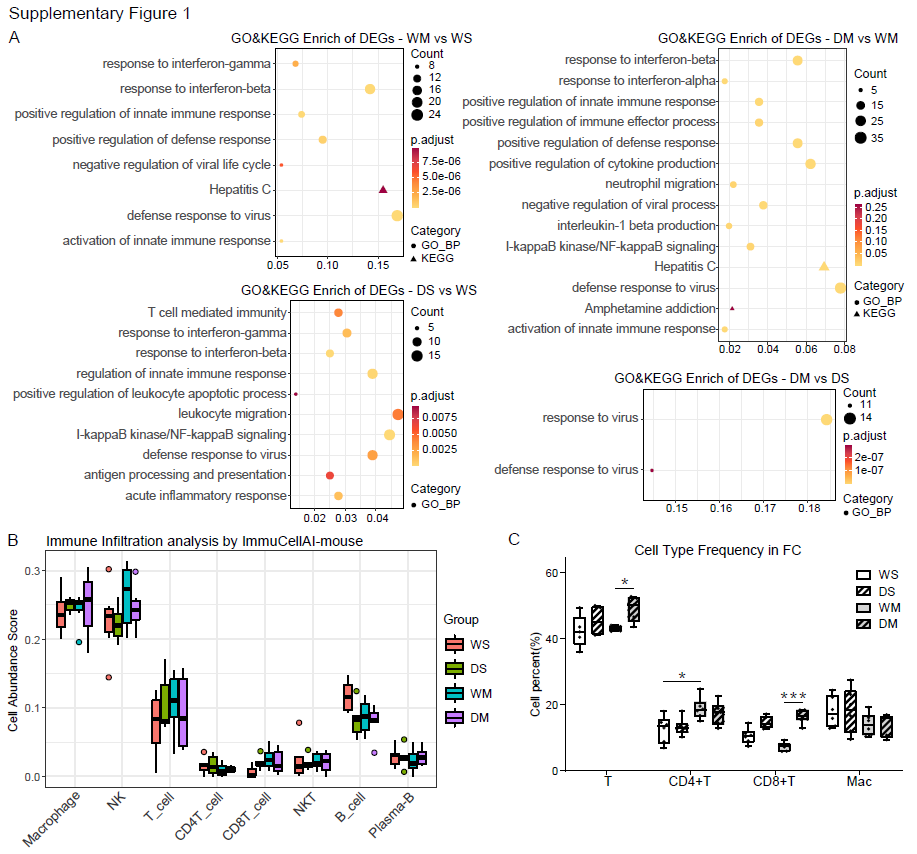

**Supplementary Figure 1.** **METH changed immune cells and immune-associated pathways in mice liver. (A)** RNA-Seq revealed changes in the immune-associated pathways in mice livers. WS: WT + Saline, n=6; WM: WT + METH, n=6; DS: DRD1 KO + Saline, n=6; DM: DRD1 KO + METH, n=6. **(B)** Immune cell abundance scores estimated by ImmuCellAI-mouse based on RNA-Seq data showed that the frequency of many immune cells may be changed after METH treatment. **(C)** Frequency of different cell types in 4 groups by flow cytometry. Group information was the same as A. *: *P* < 0.05, ***: *P* < 0.001.

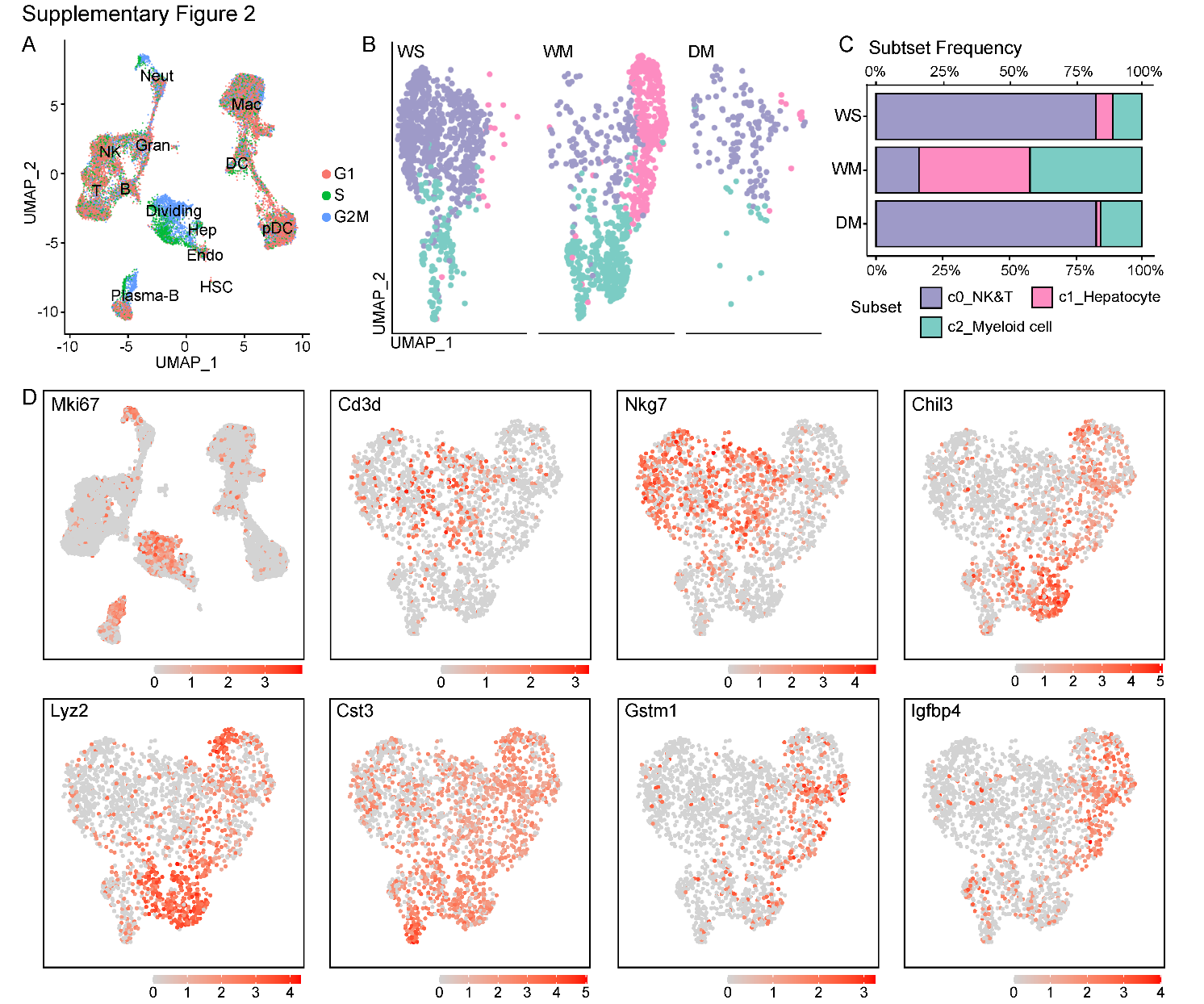

**Supplementary Figure 2.** **Dividing cell and its subclusters. (A)** The cell cycle phases for all clusters. **(B) & (C)** Group-wise cell populations and proportions of 4_Dividing cell. **(D)** The expression profiles of Mki67 in all cells and marker genes of the 4_Dividing cell population.

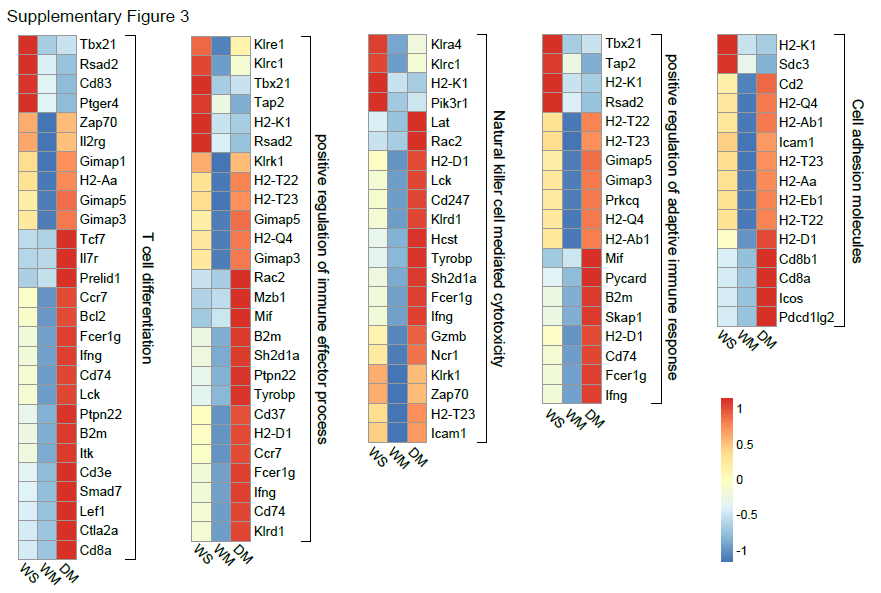

**Supplementary Figure 3. Genes changed in the representative functional pathways.** Heatmap showing the relative gene expressions in the representative functional pathways.

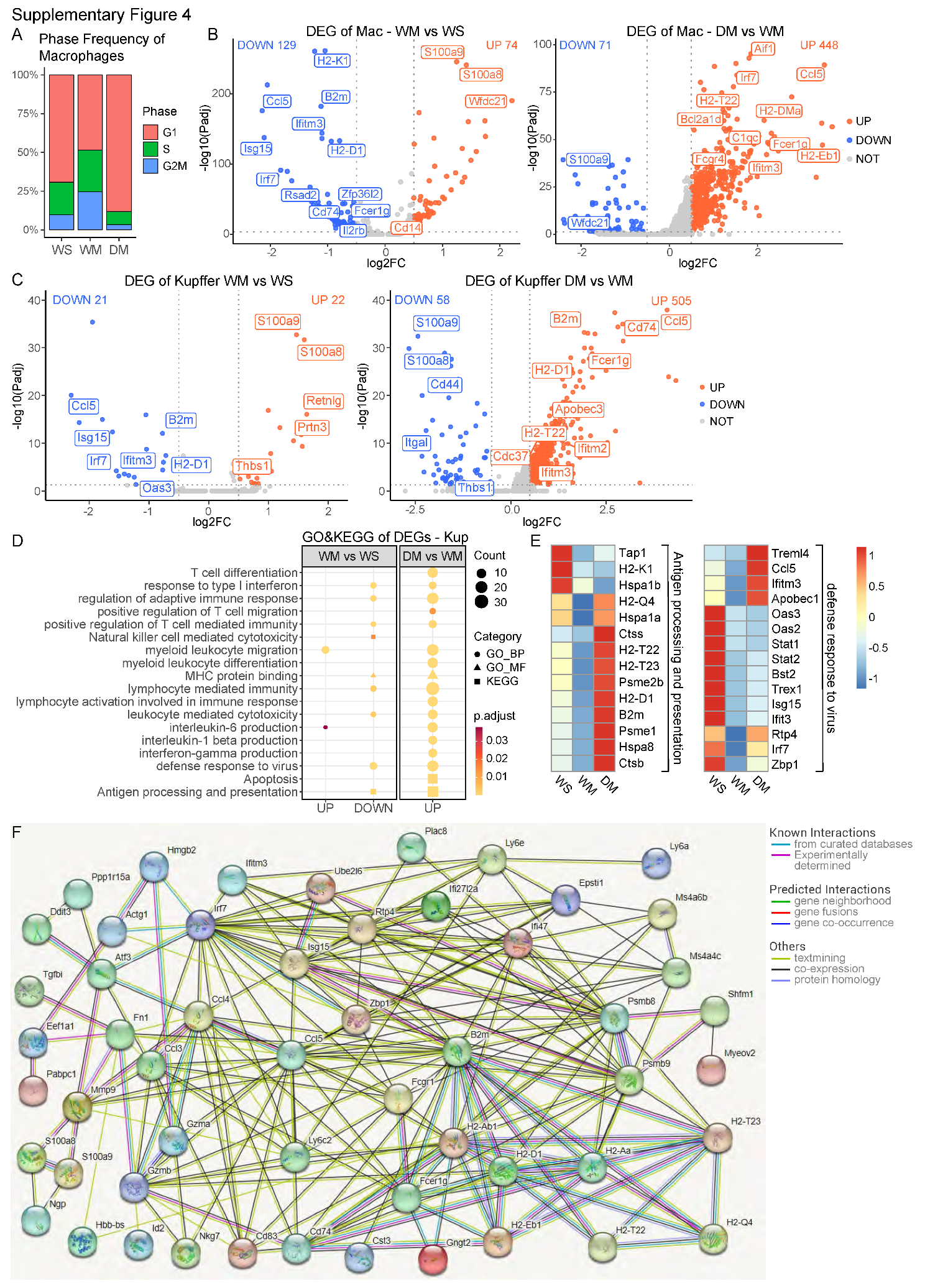

**Supplementary Figure 4. DEGs of macrophages and Kupffer cell. (A)** Cell cycle phases of macrophages from 3 groups. **(B)** Volcano plots showing fold change of gene expression (log2 scale) for down-regulated and up-regulated genes in Kupffer cells of WM vs. WS group and DM vs. WM group. Up-regulated genes (*P.adj* < 0.05 and log2FC > 0.5) are shown with red dots, down-regulated genes (*P.adj* < 0.05 and log2FC < -0.5) shown with blue dots, and insignificant genes (*P.adj* > 0.05 or log2FC < 0.5) shown with gray dots. **(C)** Volcano plots showing fold change of gene expression (log2 scale) for down-regulated and up-regulated genes in Kupffer cells of WM vs. WS group and DM vs. WM group. Up-regulated genes (*P.adj* < 0.05 and log2FC > 0.5) are shown with red dots, down-regulated genes (*P.adj* < 0.05 and log2FC < -0.5) shown with blue dots, and insignificant genes (*P.adj* > 0.05 or log2FC < 0.5) shown with gray dots. **(D)** GO and KEGG enrichment of the up-regulated and down-regulated genes of Kupffer cells in (C). **(E)** Heatmap showing the relative gene expression levels in the representative immune pathways of macrophages from 3 groups. **(F)** STRING networks of the gene sets regulated by METH through DRD1 in macrophages.

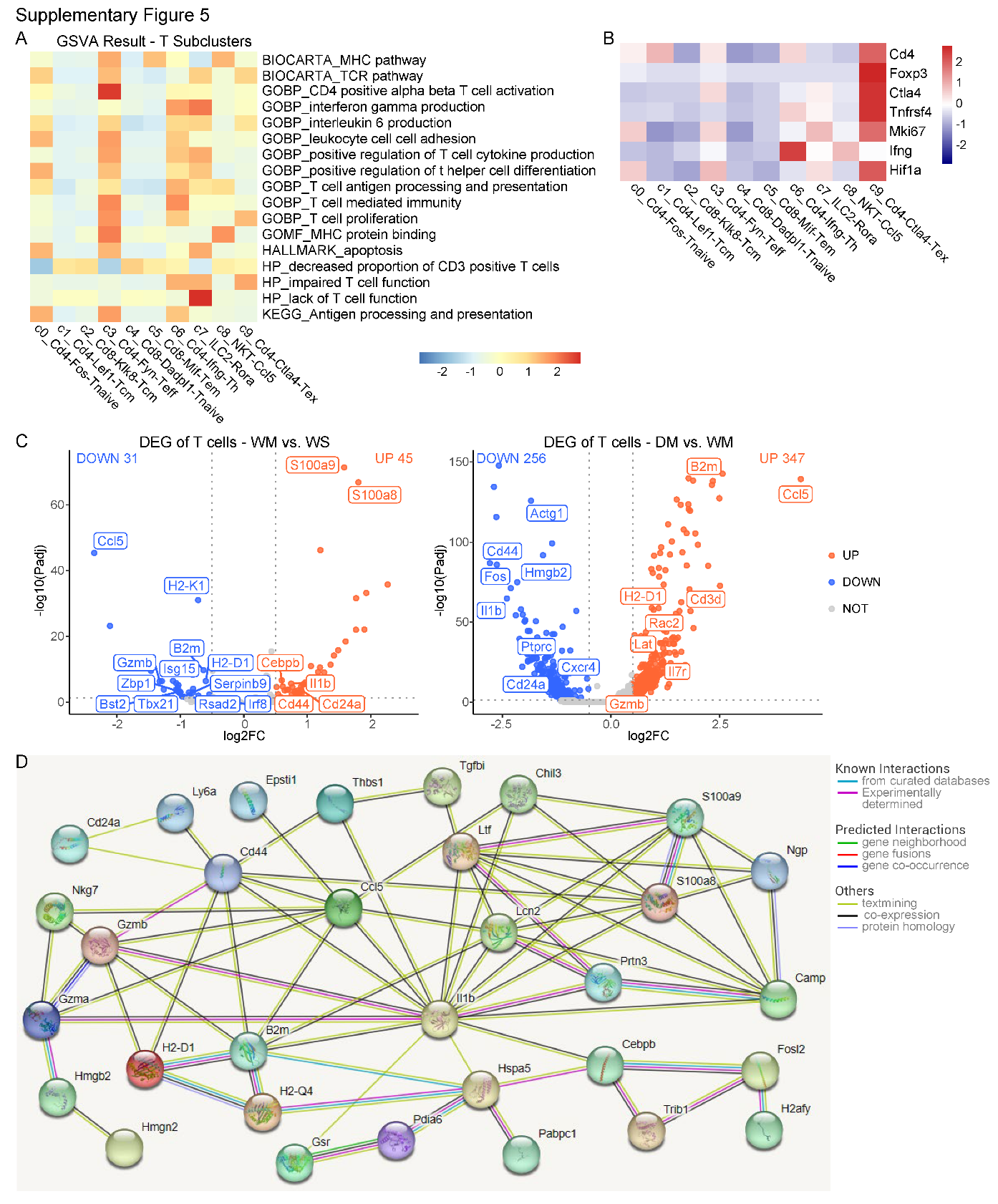

**Supplementary Figure 5. T subclusters function and DEGs. (A)** GSVA (gene set variation analysis) of T-cell subclusters. **(B)** The expressions of the feature genes from the c9_Cd4-Ctla4-Tex subcluster. **(C)** Volcano plots showing fold change of gene expression (log2 scale) for down-regulated and up-regulated genes in T cells of WM vs. WS group and DM vs. WM group. Up-regulated genes (*P.adj* < 0.05 and log2FC > 0.5) are shown with red dots, down-regulated genes (*P.adj* < 0.05 and log2FC < -0.5) shown with blue dots, and insignificant genes (*P.adj* > 0.05 or log2FC < 0.5) shown with gray dots. **(D)** STRING networks of the gene sets regulated by METH through DRD1 in T cells.

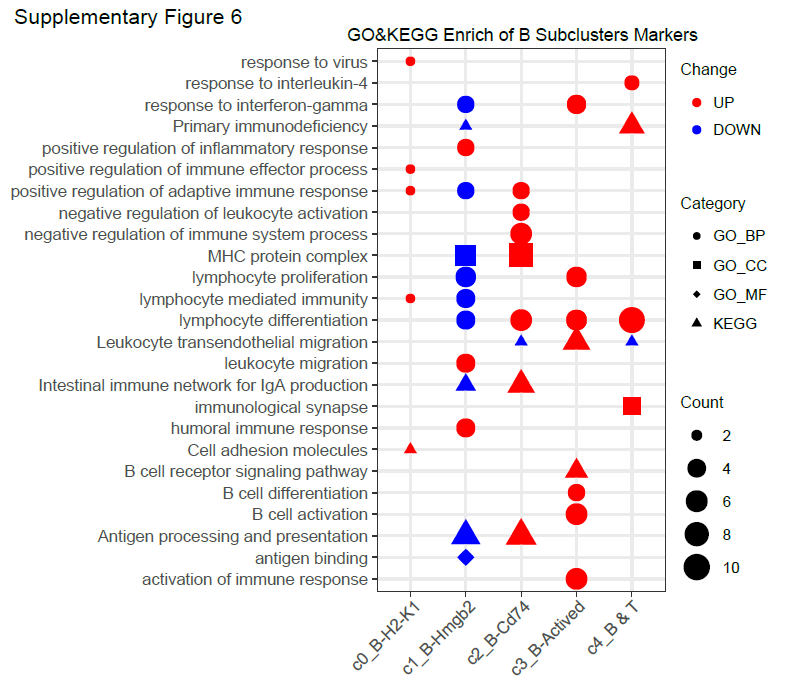

**Supplementary Figure 6. Enrichment of the marker genes of B subclusters.** GO and KEGG enrichment analysis of marker genes of five subclusters of B cells.

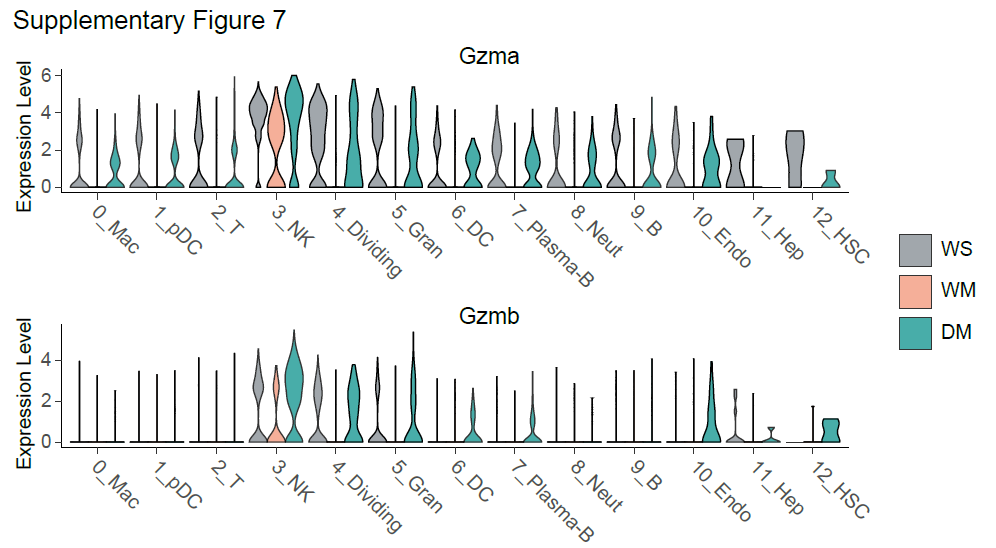

**Supplementary Figure 7. The expression levels of *Gzma* and *Gzmb*.** The expression levels of *Gzma* and *Gzmb* in all cell types from group WS, WM and DM.
